# Cellular and Molecular Resolution of Focal Segmental Glomerulosclerosis Recurrence in Human Allografts

**DOI:** 10.1101/2025.04.24.650454

**Authors:** Lorenzo Gallon, Haseeb Zubair, Thomas V. Rousselle, Amol C. Shetty, Shafquat Azim, Elissa Bardhi, Eleonora Forte, Cinthia B. Drachenberg, Enver Akalin, Manish Talwar, Jonathan S. Bromberg, Daniel G. Maluf, Valeria R. Mas

## Abstract

Primary Focal Segmental Glomerulosclerosis (FSGS) is an important cause of end-stage renal disease (ESRD). Primary FSGS recurrence rates in transplanted kidneys are high, with 25-50% in first transplants and up to 80% in second transplants, often leading to graft loss. To investigate the molecular and cellular events underlying recurrent primary FSGS (reFSGS), we performed single-nucleus RNA sequencing (snRNA-seq) on kidney transplant biopsies from patients with reFSGS and controls with normal allograft function. Our analysis revealed that podocyte loss in reFSGS is driven by metabolic and structural dysregulation rather than apoptosis. Overexpression of vascular endothelial growth factor (VEGF)-A by podocytes was observed, potentially disrupting glomerular endothelial cell growth and permeability. Parietal epithelial cells (PECs) exhibited dedifferentiation towards a podocyte-like state, potentially compensating for podocyte loss, but this was associated with increased collagen deposition and glomerular sclerosis. Ligand-receptor interactions between glomerular cells and B cells further promoted extracellular matrix deposition and fibrosis. Additionally, tubular cells demonstrated evidence of tubular sclerosis and impaired regenerative potential, accompanied by increased interactions with T cells. These findings provide novel insights into the pathogenesis of reFSGS and identify potential therapeutic targets. This study establishes a foundation for future research to further investigate cell-type-specific interventions in recurrent FSGS.

## INTRODUCTION

Primary Focal Segmental Glomerulosclerosis (FSGS) is an important cause of end-stage renal disease (ESRD) in the U.S. *(1)*, characterized by progressive scarring in the kidney’s glomeruli and the detachment of podocytes from the glomerular basement membrane (GBM). Loss of podocyte highlights glomerular dysfunction, which is a hallmark of FSGS. Despite significant advances in understanding other glomerular diseases, such as membranous nephropathy and IgA nephropathy, the mechanisms underlying primary FSGS and its recurrence following kidney transplantation remain elusive *(2–9)*. Post-kidney transplantation (KTx), primary FSGS recurs in approximately 25-50% of patients receiving their first transplant *(10–13)*, and the recurrence rate rises to 80% in subsequent transplants *(14)*. This recurrence often leads to graft loss, posing a significant challenge in the clinical management of affected patients. Current research suggests that circulating permeability factors may contribute to the disease process, but their precise role and downstream molecular mechanisms remain speculative *(15–17)*.

Recent advances in single-cell (scRNA-seq) and single-nucleus (snRNA-seq) sequencing technologies have enabled unprecedented insights into cellular heterogeneity in kidney diseases. A recent scRNA-seq propelled study identified endothelial cell-based α-2 macroglobulin (A2M) expression as a key predictor of proteinuria remission rates in FSGS of native kidneys *(18)*. Another article utilizing the similar approach demonstrated an increase in TNF signaling in the tubular region of kidneys with FSGS *(19)*. ScRNAseq of urinary samples from FSGS patients identified immune cells, mainly monocytes, and renal epithelial cells, with shed podocytes showing high expression of epithelial-to-mesenchymal transition (EMT) markers *(20)*. However, studies focusing specifically on recurrent FSGS and even other glomerulopathies in transplant recipients are lacking. Leveraging snRNA-seq, which preserves the integrity of cellular nuclei in archived biopsy samples, offers a unique opportunity to interrogate the molecular and cellular underpinnings of recurrent FSGS at high resolution *(21, 22)*.

In this study, we demonstrate the use snRNA-seq on kidney biopsy samples from transplant recipients with recurrent FSGS (reFSGS) and controls with normal allograft function. Analysis of the transcriptomic data we identified key alterations in podocyte structure, PEC differentiation, and tubular cell function that contribute to reFSGS pathogenesis. These findings establish a framework for understanding disease recurrence and highlight potential therapeutic targets for intervention.

## RESULTS

### Comprehensive identification of kidney cell types through snRNAseq

The characteristics of patients and samples used in this study are shown in **Table 1**, and the histological findings are reported in **Table 2**. Quality control results are provided in **Fig. S1** and **Tables S1 and S2**. Thirteen human kidney tissue needle biopsies, **Fig. 1A**, yielded 64,694 nuclei: these were composed of three normal native kidneys (nNK; 12,993 nuclei; nuclei from GSE131882 *(23)*), one native kidney with FSGS (nFSGS; 5,546 nuclei), four normal allografts (NKTx; 27,957 nuclei), three kidney allografts at the time of recurrent FSGS (KTxFSGS-T1; 12,361 nuclei), and two follow-up collections from these patients (KTxFSGS-T2; 5,837 nuclei). An integrated UMAP generated 19 clusters, **Figs. 1B and S2**, wherein we identified podocytes (POD), parietal epithelial cells (PEC), endothelial cells (EC), tubule cells and collecting duct cells, immune (IMM) cells, fibroblasts (FIB), mesangial (MES) cells, myocytes (MYO), and two uncharacterized cell clusters. Details of cell markers (Fold change, FC, > 1.5; adjusted (adj.) *p*-value < 0.01) and nuclei per cluster are in **Tables S3 and S4**. Clusters with fewer than five marker genes were excluded as over-clustering artifacts. The proportions of cell clusters under each pathological condition and their marker profiles are shown in **Figs. 1C and D**.

**Fig. 1:**
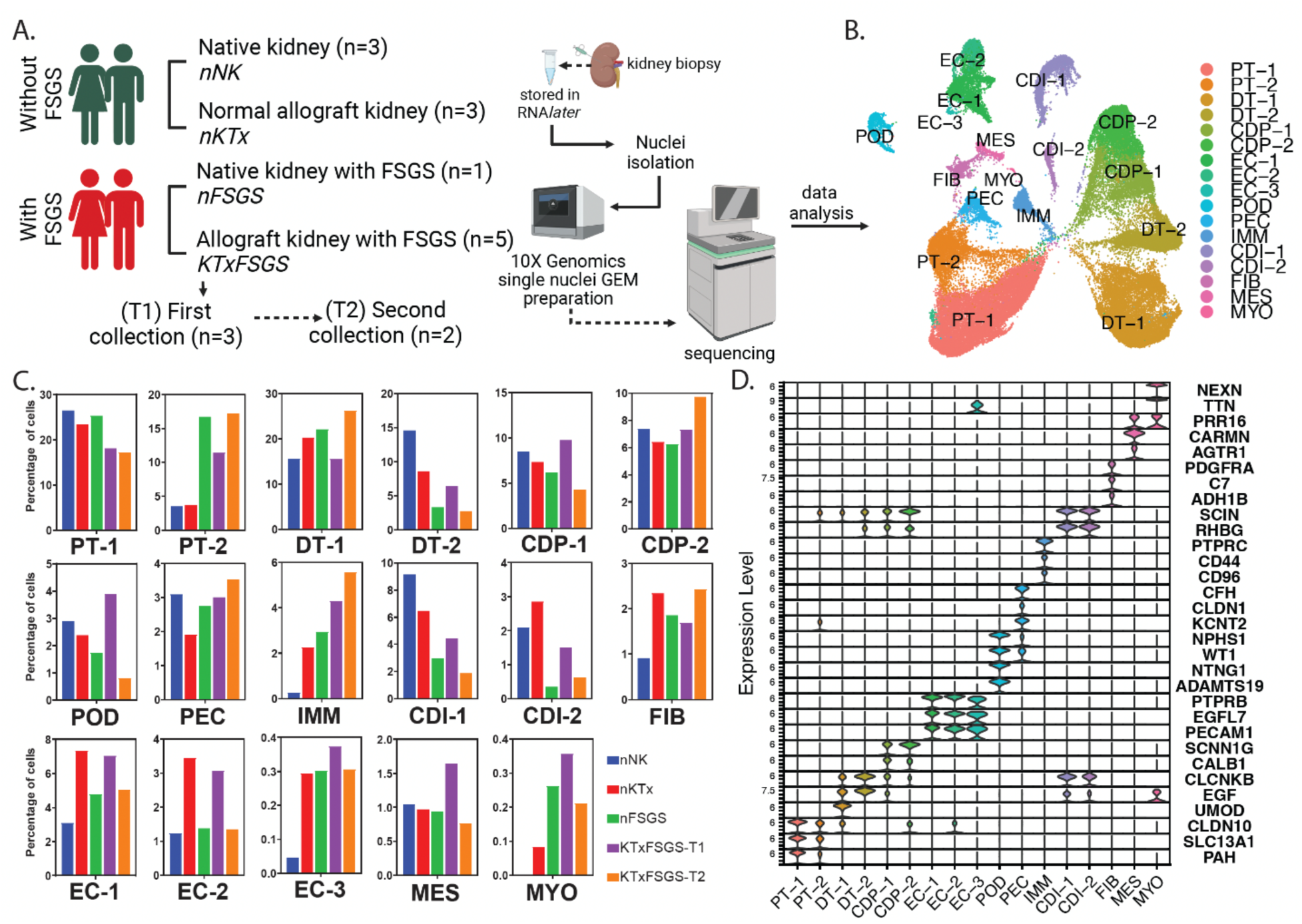
Study design and characterization of main cell clusters. (A) Study workflow. (B) Cluster analysis identified seventeen major cell types. (C and D) Cell proportions and major cell markers used to identify cell clusters. Cell clusters include podocytes (POD), parietal epithelial (PEC), endothelial (EC-1, EC-2, EC-3), proximal tubule (PT-1 & PT-2), distal tubule (DT-1 & DT-2), collecting duct principal (CDP-1 & CDP-2), collecting duct intercalated (CDI-1 & CDI-2), immune (IMM), fibroblast (FB), mesangial (MES), myocyte (MYO).

**Table 1:** Characteristics of patients and donors from samples included in the study.

| Condition | Sample ID | ID | Donor Characteristics |  |  | Recipient Characteristics |  |  |  |  |
| --- | --- | --- | --- | --- | --- | --- | --- | --- | --- | --- |
|  |  |  | Age | Race | Sex | Age | Race | Sex | sCR* | Proteinuria* |
| Native Kidneys (nNK) | GEO Dataset | GSM3823 939^ | 54 | C | M | -- | -- | -- | 1.28 | -- |
|  | GEO Dataset | GSM3823 940^ | 62 | C | M | -- | -- | -- | 1.21 | -- |
|  | GEO Dataset | GSM3823 941^ | 61 | C | F | -- | -- | -- | 0.89 | -- |
| Normal Allograft (NKTx) | KUT-023 | NKTx1 | 21 | C | M | 70 | AA | M | 0.80 | none |
|  | KUT-060 | NKTx2 | 49 | NA | M | 56 | AA | F | 1.00 | none |
|  | KUT-081 | NKTx3 | 17 | C | M | 32 | C | F | 0.76 | none |
|  | NW-4 | NKTx4 | 23 | H | F | 46 | C | F | 1.00 | none |
| Native Kidney FSGS (nFSGS) | NW-3 | nFSGS | 63 | C | F | -- | -- | -- | 2.09 | 4.0 |
| Post-KTx-FSGS (KTx-FSGS) | NW-1 | T1-FSGS1** | 49 | C | F | 64 | C | M | 1.95 | 7.6 |
|  | NW-6 | T1-FSGS2** | 40 | C | M | 65 | AA | M | 2.2 | 4.8 |
|  | NW-5 | T1-FSGS3 | 52 | C | M | 37 | C | M | 4.0 | 4.0 |
|  | NW-2 | T2-FSGS1** | Refer T1-FSGS#1 |  |  |  |  |  | 2.18 | 3.8 |
|  | NW-7 | T2-FSGS2** | Refer T1-FSGS#2 |  |  |  |  |  | 1.6 | 4.0 |

NA, Not available; C, Caucasian; AA, African American; H, Hispanic; F, Female; M, Male; FSGS, Primary Focal Segmental Glomerular Sclerosis; T1: first recurrence, T2: second recurrence. Age is denoted in years; sCR, serum creatinine levels denoted in mg/dL; proteinuria in g/day (is it spot urine protein/creatinine ratio or 24 hour urine collection?); time of collection denoted in months; \*: donor sCR and proteinuria at time of biopsy; \*\*: longitudinal paired biopsies (T1-FSGS1 and T2-FSGS1; and T1-FSGS2 and T2-FSGS2); ^: samples accessed through GEO Series accession number GSE131882.

**Table 2:**
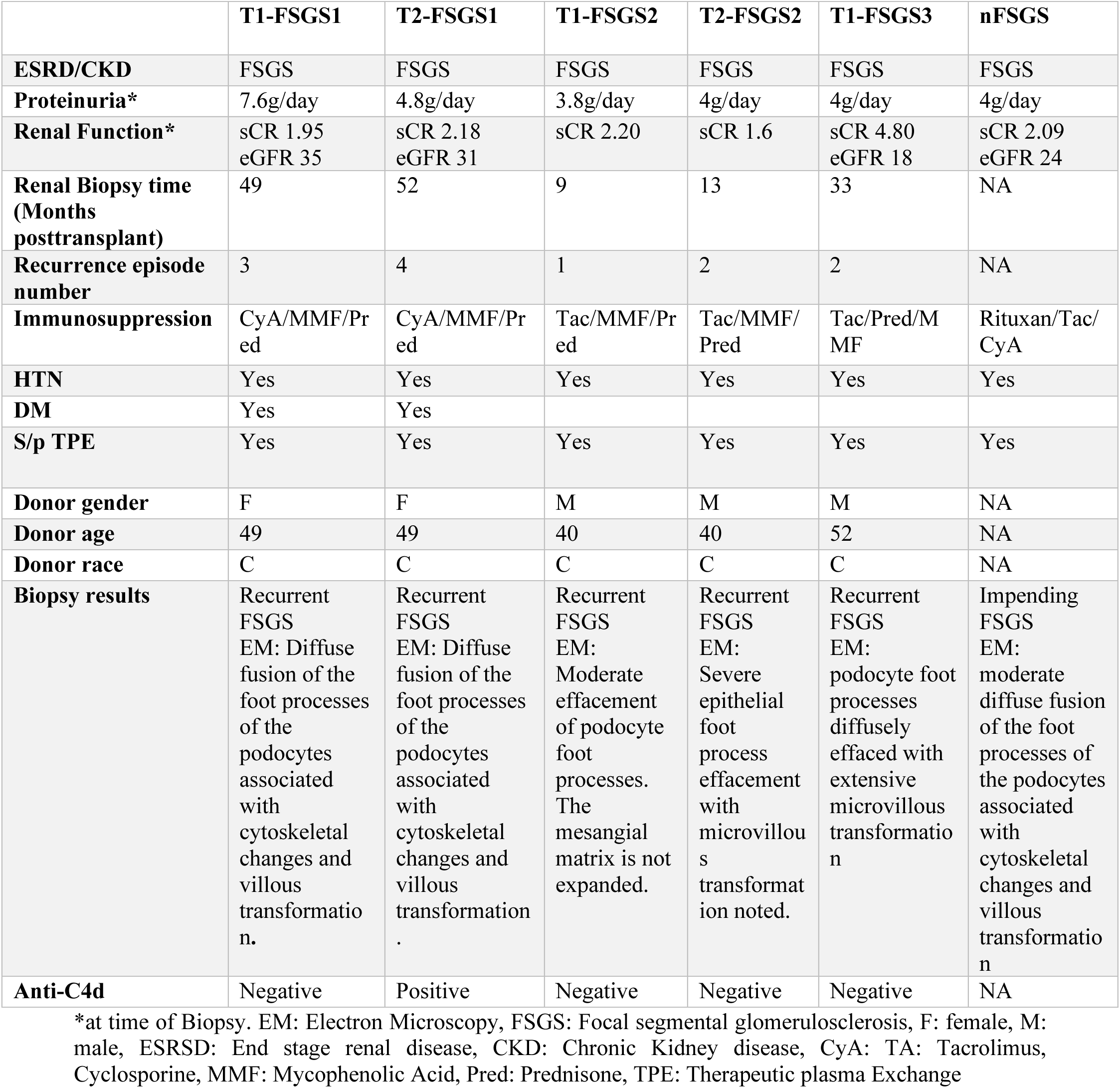
Clinical and histological evaluations of biopsy samples from patients with FSGS.

|  | **T1-FSGS1** | **T2-FSGS1** | **T1-FSGS2** | **T2-FSGS2** | **T1-FSGS3** | **nFSGS** |
| --- | --- | --- | --- | --- | --- | --- |
| **ESRD/CKD** | FSGS | FSGS | FSGS | FSGS | FSGS | FSGS |
| **Proteinuria\*** | 7.6g/day | 4.8g/day | 3.8g/day | 4g/day | 4g/day | 4g/day |
| **Renal Function\*** | sCR 1.95<br>eGFR 35 | sCR 2.18<br>eGFR 31 | sCR 2.20 | sCR 1.6 | sCR 4.80<br>eGFR 18 | sCR 2.09<br>eGFR 24 |
| **Renal Biopsy time (Months posttransplant)** | 49 | 52 | 9 | 13 | 33 | NA |
| **Recurrence episode number** | 3 | 4 | 1 | 2 | 2 | NA |
| **Immunosuppression** | CyA/MMF/Pred | CyA/MMF/Pred | Tac/MMF/Pred | Tac/MMF/Pred | Tac/Pred/MMF | Rituxan/Tac/CyA |
| **HTN** | Yes | Yes | Yes | Yes | Yes | Yes |
| **DM** | Yes | Yes |  |  |  |  |
| **S/p TPE** | Yes | Yes | Yes | Yes | Yes | Yes |
| **Donor gender** | F | F | M | M | M | NA |
| **Donor age** | 49 | 49 | 40 | 40 | 52 | NA |
| **Donor race** | C | C | C | C | C | NA |
| **Biopsy results** | Recurrent FSGS<br>EM: Diffuse fusion of the foot processes of the podocytes associated with cytoskeletal changes and villous transformation. | Recurrent FSGS<br>EM: Diffuse fusion of the foot processes of the podocytes associated with cytoskeletal changes and villous transformation. | Recurrent FSGS<br>EM: Moderate effacement of podocyte foot processes. The mesangial matrix is not expanded. | Recurrent FSGS<br>EM: Severe epithelial foot process effacement with microvillous transformation noted. | Recurrent FSGS<br>EM: podocyte foot processes diffusely effaced with extensive microvillous transformation | Impending FSGS<br>EM: moderate diffuse fusion of the foot processes of the podocytes associated with cytoskeletal changes and villous transformation |
| **Anti-C4d** | Negative | Positive | Negative | Negative | Negative | NA |

\*at time of Biopsy. EM: Electron Microscopy, FSGS: Focal segmental glomerulosclerosis, F: female, M: male, ESRD: End stage renal disease, CKD: Chronic Kidney disease, CyA: Tacrolimus, Cyclosporine, MMF: Mycophenolic Acid, Pred: Prednisone, TPE: Therapeutic plasma Exchange

### Podocyte transcriptional changes highlight cytoskeletal dynamics in recurrent FSGS post-kidney transplant

Further characterization of podocytes (POD), parietal epithelial cells (PEC), and mesangial cells (MES) —the principal glomerular cells—is presented in **Figs. 2A and B**. Some of the marker genes defining PODs had an overlap with PECs and MES (adj. *p*-value < 0.01, **Table S5, Fig. 2B**). However, podocyte-specific essential (*NPHS1*, *NPHS2*, *WT1*, *PODXL*, *LAMB2*, *PLCE1*) and cytoskeletal markers (*ACTN4*, *SYNPO, MYH9, CD2AP, DST*, *EZR*) were uniquely present in this cluster (**Fig. 2C**) *(24)*. A trend of reduction in podocyte proportions in KTxFSGS overtime was observed. Specifically, the percentage of podocytes was 6.0%, 0.9%, and 1.9% in samples from three different recurrent patients collected at 9-, 33-, and 49-months (mo) post-KTx, respectively (**Table S3**). In the follow-up samples (KTxFSGS-T2), the percentage of recovered podocytes decreased from 6.0% to 2.6%, and from 1.9% to 0.7%, respectively. Indeed, nFSGS too had a lower proportion of podocytes relative to the nNK. Gene expression analysis between KTxFSGS-T1 to NKTx identified 141 downregulated and 8 upregulated genes (adj. *p*-value < 0.05, |fold change| > 1.5, **Fig. S3**, **Table S6**). Genes downregulated included in the POD cluster during FSGS recurrence and included *STAT1*, *NFKB1*, *LRP6*, *ROBO1*, *NPHS2*, *MALAT1*, *EGF*, *ERBB4*, *BACH1,* and had an upregulation of *VEGFA* (**Fig. 2D**) *(25–32)*. Interestingly, *in vitro* treatment of conditionally immortalized podocytes (c-AP Podocytes) with serum collected from patients at the time of the first biopsy collection (KTxFSGS-T1) reduced the expression of multiple genes including *LAMB1, CD44, WT1*, and a patient-dependent but reduction in expression of *STAT1*, *MALAT1* **(Fig. S3)**, potentially reflecting the heterogeneity in expression of the ‘causal’ factor in the serum. Consistent with our single nucleus RNA-seq findings where we observed a reduced *EGF* expression in podocytes, immunohistochemistry demonstrated markedly decreased and diffused EGF protein staining within glomerular regions of diseased kidneys. (**Fig. 2E**).

**Fig. 2:**
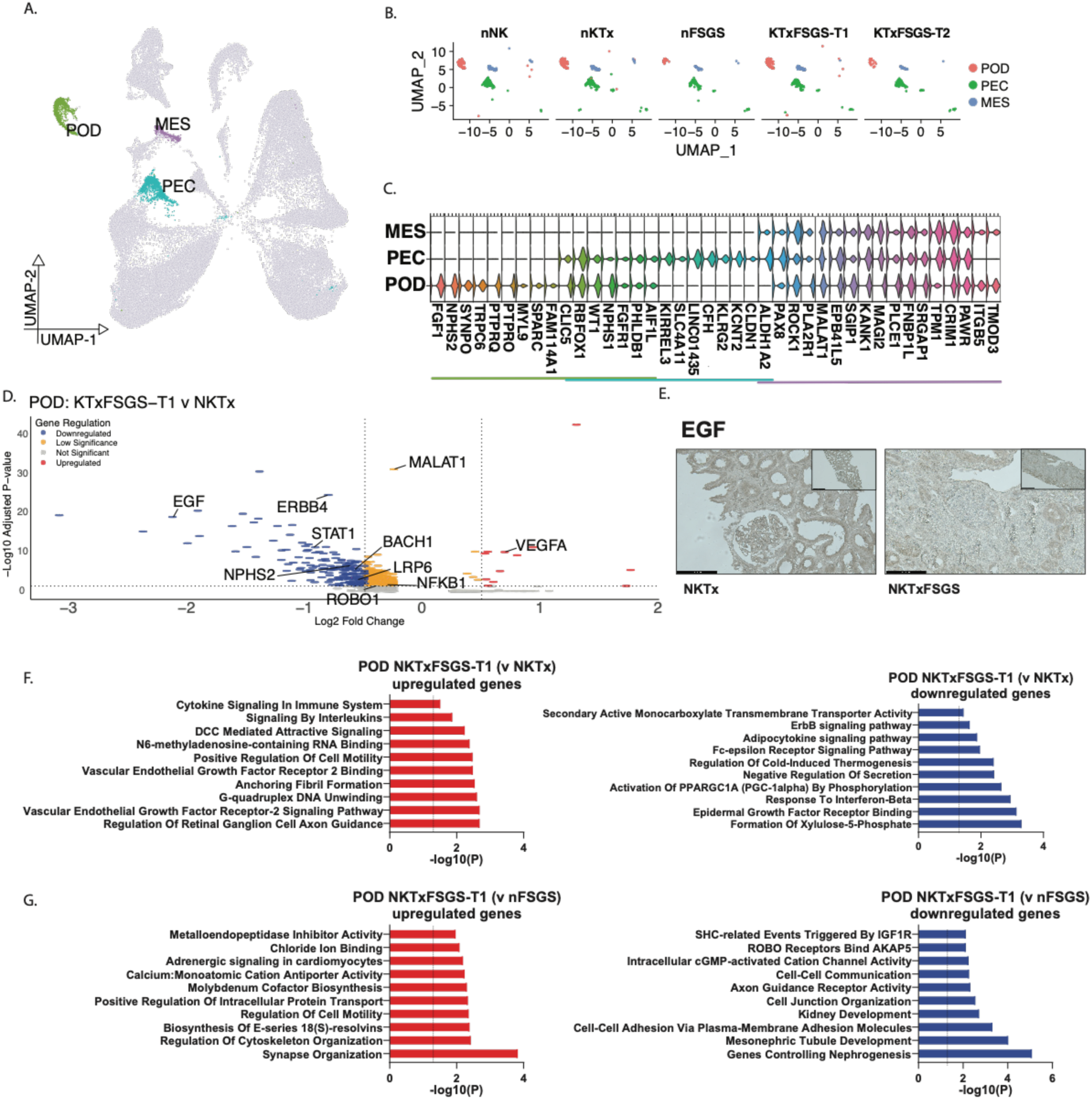
Molecular and cellular properties of glomerular cells. (A and B) UMAP identifies the podocytes (POD), parietal epithelial cells (PEC), and mesangial cells (MES) among the different conditions. (C) Unique and overlapping gene markers in POD, PEC, and MES cell clusters. (D) Split violin plots for transcription factors and structural gene expression between NKTx and KTxFSGS-T1. (E) Immunohistochemical analysis of EGF expression from NKTx and KTxFSGS biopsies. Gene ontology using differentially expressed genes between (F) KTxFSGS-T1 v. NKTx, and (G) KTxFSGS-T1 v. nFSGS.

A detailed examination of the downregulated genes revealed reduction of multiple Phospholipase C (PLC) enzyme; *PLCE1*, *PLCL1*, *PLCG2*. We also observed a significant downregulation of two calcium-permeable channels, *TRPC1* and *TRPM1*. Additionally, we observed dysregulation in carbohydrate metabolism, potentially through the development of insulin resistance, indicated by the downregulation of *INSR*. The downregulated genes were also linked to a reduction in IL-1 signaling and IFN-gamma response. There was also a decrease in ERBB signaling in these podocytes due to the decrease in *EGF* and *ERBB4* expression. While overactivation of the ERBB/EGFR signaling pathways has been reported to induce injury *(33, 34)*, EGFR pathway has a crucial role in repair of damaged kidneys *(35)*. Since these signaling pathways are crucial for cell survival, differentiation, and growth, their downregulation implies compromised podocyte viability and functionality, potentially contributing to the pathological process of FSGS recurrence (**Fig. 2F**). GO biological process analysis of over-expressed genes in POD from KTxFSGS-T1 samples identified multiple processes/pathways involving *VEGFA*. Overexpression of *VEGFA* in podocytes suggests a potential compensatory response to hypoxic conditions and is associated with induced endothelial injury *(36, 37)*. In podocytes, VEGF-A overexpression has been demonstrated to increase VEGFR2 phosphorylation, leading to proteinuria due to increased glomerular hypertrophy *(38–40)*. Gene expression profiles of podocytes from KTxFSGS-T1 with those from nFSGS and identified 34 differentially expressed genes (DEGs), indicating minimal differences in podocytes between kidney grafts and native kidneys with FSGS. The DEGs were associated with pathways involved in cell-cell adhesion, cell-cell communication, and cell junction organization, and were downregulated in recurrent FSGS **(Fig. 2G)**. Two podocyte-injury relevant upregulated genes in KTxFSGS-T1 podocytes included *HPGD* (log_2_FC of 6.5 with adj. p-value of 2x10^-04^) and *SLC8A1* (log_2_FC of 2.4 with adj. p-value of 1.72x10^-6^) (**Table S7**). *HPGD* upregulation in podocytes may modulate inflammatory responses, as it is involved in prostaglandin metabolism. Aiming to evaluate the extent of metabolic dysfunction, we reevaluated the DEGs with a lesser stringent filtering (adj. p-value < 0.05 alone, **Fig. S4A, Table S6**) resulting in the identification of which resulted in 450 downregulated, and 16 upregulated genes in the KTxFSGS-T1 group. This approach allowed for a more detailed identification of significantly altered genes associated with relevant pathways. We focused on fatty acid metabolism pathways and observed an enrichment of transcription factor genes associated with adipogenesis (**Fig. S4B**). Additionally, there was a notable reduction in leptin signaling, prostaglandin synthesis, and lipolysis regulation in adipocytes. In addition to these, the downregulated genes also associated with reduction in cytoskeleton organization, positive regulation of carbohydrate metabolic process, and the regulation of plasma membrane bounded cell projection assembly. These factors lead to podocyte detachment and eventual urinary excretion rather than the activation of cell death pathways.

### Decoding temporal variations in podocyte behavior in post-transplant FSGS recurrence

Although the three graft biopsies of KTxFSGS-T1 had similar histological characteristics, they were collected at different times, likely reflecting various disease stages. Thus, we examined how these temporal differences appeared at the transcriptional level in podocyte clusters from each KTxFSGS-T1 patient sample to determine if unique podocyte transcriptional profiles could capture differences in disease progression. We compared gene expression profiles of podocytes from patient T1-FSGS1 (49 months (mo) post-KT, third FSGS recurrence) with those from NKTx, identifying 55 DEGs (**Table S8**). Downregulated genes were linked to kidney development and various metabolic pathways (carbohydrate, fatty acid metabolism, and lipolysis regulation). Upregulated genes were related to nephrogenesis, migration, and protein localization to membrane (**Fig. S5A**). In the T1-FSGS2 sample (9 mo post-KT, first FSGS recurrence), we found a smaller number of DEGs (20), mostly upregulated (n=17, **Table S9**). Notably, CD44 (Log_2_FC = 3.4, adj. p-value = 4.16e-13) was significantly increased in the POD cluster, expressed by 28% of cells in the FSGS2 sample (pct1: 0.28, **Fig. S6**). CD44 upregulation, common in FSGS and other glomerulopathies, may be essential for podocyte survival or regeneration *(41–44)*. Comparing T1-FSGS3 (33 mo post-KT, second recurrence) with NKTx, we identified 167 DEGs, mostly downregulated in T1-FSGS3 (**Table S10**). Downregulated genes in our analysis were associated with protein localization, cell-cell adhesion, cyclic nucleotide catabolism, prostaglandin, and leukotriene metabolism in senescence (**Fig. S5B**).

### Parietal epithelial cells transcriptional profile identifies subsets with potential regenerative capacity

The PEC cluster (n = 1,539 cells) exhibited enriched expression of *CLDN1*, an established PEC marker (**Fig. 1D**). Intriguingly, it also showed markers for POD (*WT1, NPHS1*) and progenitors (*PROM1;* **Fig. 2B, 3A**, and **Table S5**). Notably, 40% of PECs expressed *PROM1* (**Fig. 3B**). We identified a distinct PEC subset, previously known as *parietal podocytes (45)*, *WT1*+ PECs co-expressing *NPHS1*, which can transition toward the podocyte lineage (**Fig. 3A** and **B**). A relative decrease in these WT1+ (*PROM1*+ or *PROM1*-) PECs was observed in patients with FSGS, both in native and graft kidney samples. Upregulated genes in PEC *WT1*+ *PROM1*+ showed enriched terms related to renal system development and cell morphogenesis compared to PEC *WT1*+ *PROM1*-. There was a correlation between the loss of POD and decreased PEC differentiation in FSGS (**Fig. 3B–inset**). This reduction in dedifferentiation was accompanied by an increase in PECs in the S cell cycle phase in individual KTxFSGS-T1 cases, indicating PEC proliferation (**Fig. 3C)**.

**Fig. 3:**
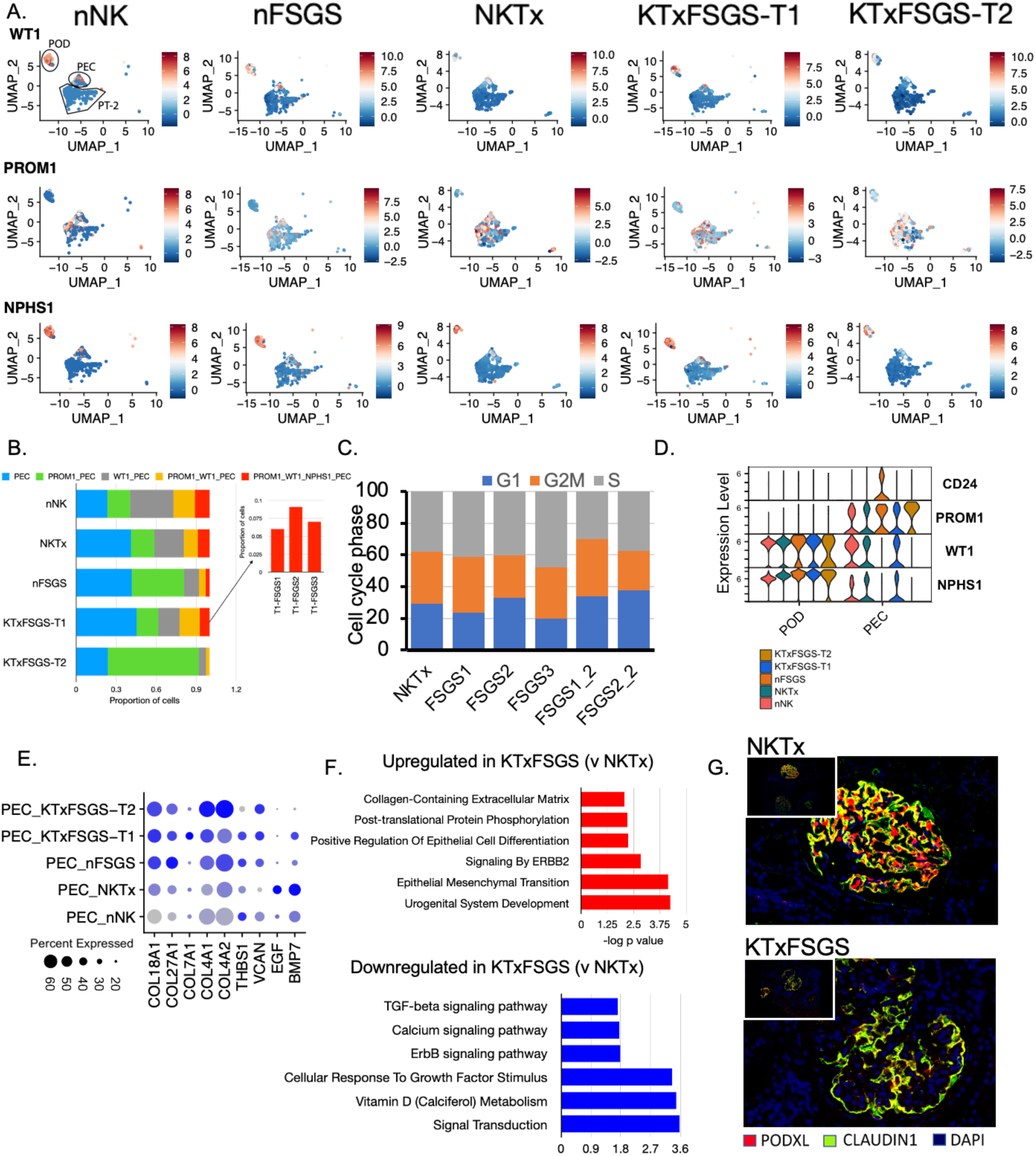
The PEC subcluster analysis. (A) The expression of podocyte marker, *WT1* and *NPHS1*, and regeneration and repair marker, *PROM1*, was determined in the three clusters, POD, PEC and PT2. (B) Percentage of cells with expression of *WT1*, *PROM1, NPHS1*. (C) Cell cycle status of PEC based on per KTxFSGS-T1 sample. (D) High expression of ECM genes as observed in KTxFSGS-T1. (E) Expression of *PROM1*, *CD24, WT1, NPHS1,* and *PODXL* in different conditions. (F) Pathway analysis of DEGs between KTxFSGS-T1 v. NKTx. (G) Immunofluorescence staining of NKTx and KTxFSGS for PODXL and Claudin-1, a representative glomerulus has been shown.

We performed a comprehensive analysis of the expression levels of *PODXL*, *PROM1*, *WT1*, and *NPHS1* in POD and PECs across different patient groups. Given recent literature suggesting the co-expression of *PROM1* and *CD24* on the surface of PEC progenitor cells located at the tubular pole *(46, 47)*, we also included CD24 in our examination. Notably, *CD24* expression was observed exclusively in nFSGS and was absent in KTxFSGS (**Fig. 3D**). Furthermore, there was an increase in the expression of ECM genes such as *COL18A1, COL27A1, COL7A1, COL4A1, COL4A2, THBS1, and VCAN* in KTx-FSGS PECs compared to NKTx PECs (**Fig. 3E**). However, no difference in ECM-related genes was observed between nFSGS and KTxFSGS PECs. CD44 is a marker of activated PECs, which are stimulated by ligands such as HB-EGF. Activated PECs are known to contribute to glomerular injury and fibrosis. In this study, *CD44* expression was absent in PECs from nNK, NKTx, and nFSGS. However, there was a relative increase in *CD44* and *HB-EGF* expression in PECs from recurrent FSGS (KTx-FSGS-T1) compared to NKTx, indicating PEC activation in recurrent FSGS (**Fig. S7A**).

Pathway enrichment analysis revealed that downregulated genes in PECs from KTxFSGS-T1 compared to NKTx are involved in vitamin D metabolism, nuclear receptor signaling, signal transduction, and cellular response to growth factor stimuli (**Fig. 3F**). Notably, the downregulated genes included orphan nuclear receptors (*ESRRG, RORA*, and *NR3C2*), with RORA and NR3C2 playing roles in cell differentiation (**Fig. S7B**, **Table S6**). *RORA* has been observed to regulate cell differentiation in HIV-associated nephropathy *(48)*. Upregulated genes in PECs from KTxFSGS-T1 were associated with activation of ERBB2, post-translational protein phosphorylation, positive regulation of epithelial cell differentiation involved in kidney development, and collagen-containing ECM, suggesting increased ECM deposition. Importantly, representative immunohistochemistry staining showed increased co-expression of PODXL (a podocyte marker) and Claudin-1 (a PEC marker), with minimal “podocyte only” expression of PODXL (**Fig. 3G**). This indicates a breakdown in the distinct roles of podocytes and PECs. The observed staining patterns suggest potential proliferation of epithelial cells without proper differentiation, contributing to glomerulosclerosis and functional impairment in FSGS. The disorganization and disruption in staining patterns reflect the underlying damage to the glomerulus and the progression of the disease. It is important to note that the expression of Claudin-1 in the lining of the Bowman’s capsule as well as in the glomerular cells suggests a mild, subclinical response to the transplantation process, suggesting that PECs are actively involved in maintaining and normalizing glomerular integrity following minor injuries. The activation of PECs to podocytes, impaired differentiation, and ECM deposition are key contributors to glomerular scarring in recurrent FSGS. These findings highlight the critical role of PEC activation and impaired differentiation in the progression of recurrent FSGS, with significant implications for understanding the mechanisms of glomerular injury and fibrosis in this condition.

### Endothelial cell dysfunction in FSGS

Three unique clusters of ECs were identified in our dataset (**Fig. 4A**). The cell markers for the three major EC cell sub-clusters demonstrated the presence of **glomerular** (Cluster EC-2; *PECAM1*, *KDR*, *EHD3*), **peritubular** (Cluster EC-1; *PECAM1*, *KDR^-^*), and **lymphatic** (Cluster EC-3; *PECAM1*, *KDR*, *PROX1*, *LYVE1*) endothelial cell types of renal system (**Fig. 4B**)*(49, 50)*. The proportions of peritubular ECs were reduced in nFSGS compared to nNK and both peritubular and lymphatic ECs were reduced in KTxFSGS-T1 compared to NKTx, respectively Conversely, an increase in glomerular ECs was observed in both native and allograft FSGS (**Fig. 4C**). This shift in EC populations indicates significant vascular changes associated with native FSGS and its recurrence post KTx. Considering the glomerular nature of the disease, we focused on identifying molecular changes in nFSGS and KTxFSGS and their effect on the glomerular EC. Compared to nNK, nFSGS showed a reduction in the expression of *NRXN1*. This gene codes for a membrane protein of the neurexin family, which is primarily present on neuronal cells but also found on ECs, where it plays a role in angiogenesis *(51)*. Indeed, there was an enrichment of pathways associated with vasculogenesis and tissue remodeling (**Table S11)** due to the upregulation of *KDR, TIE1,* and *TGFB1*. The upregulation of *KDR* (which codes for the protein VEGFR2) aligns with the increased *VEGFA* observed in podocytes. This explains the increased proportion of glomerular ECs (**Fig. 4C**). This interaction of VEGF-A (upregulated in FSGS-associated podocytes) with VEGFR2 induces angiogenesis *(40, 52)*. Additionally, there was enrichment of Wnt signaling pathways, which are also known to cause angiogenesis *(53)*. Some pathways observed to be downregulated in nFSGS are associated with negative regulation of cellular component movement, Fc gamma R-mediated phagocytosis, and fluid shear stress and atherosclerosis (**Fig. S8**). These downregulated pathways indicate a potential impairment in cellular mobility, immune response mechanisms, and vascular integrity in the glomeruli. In KTxFSGS, there was reduced expression of genes responsible for ion transport and hormone-mediated signaling pathways, and metabolism regulation (**Fig. 4D, Table S12**). The imbalance resulting from alteration in sodium and chloride ion transporters can result in increased extracellular sodium and thereby increase stiffness of the endothelial cells, weakening the endothelial glycocalyx leading to higher permeability *(54, 55)*. Additionally, the downregulation of *PPARA* in endothelial cells can alter the repair process of damaged endothelial cells *(56)*. An interesting observation from the EC DEGs was a correlation between phosphodiesterase, *PDE10A*, *PDE4D*, and *PDE1A* (**Fig. 4E**). The activity of phosphodiesterase is hypothesized to maintain EC permeability, and their loss can lead to altered permeability *(57–59)*. Thus, altered molecular events during KTxFSGS are associated with disruptions in metabolism and endothelial cell permeability.

**Fig. 4:**
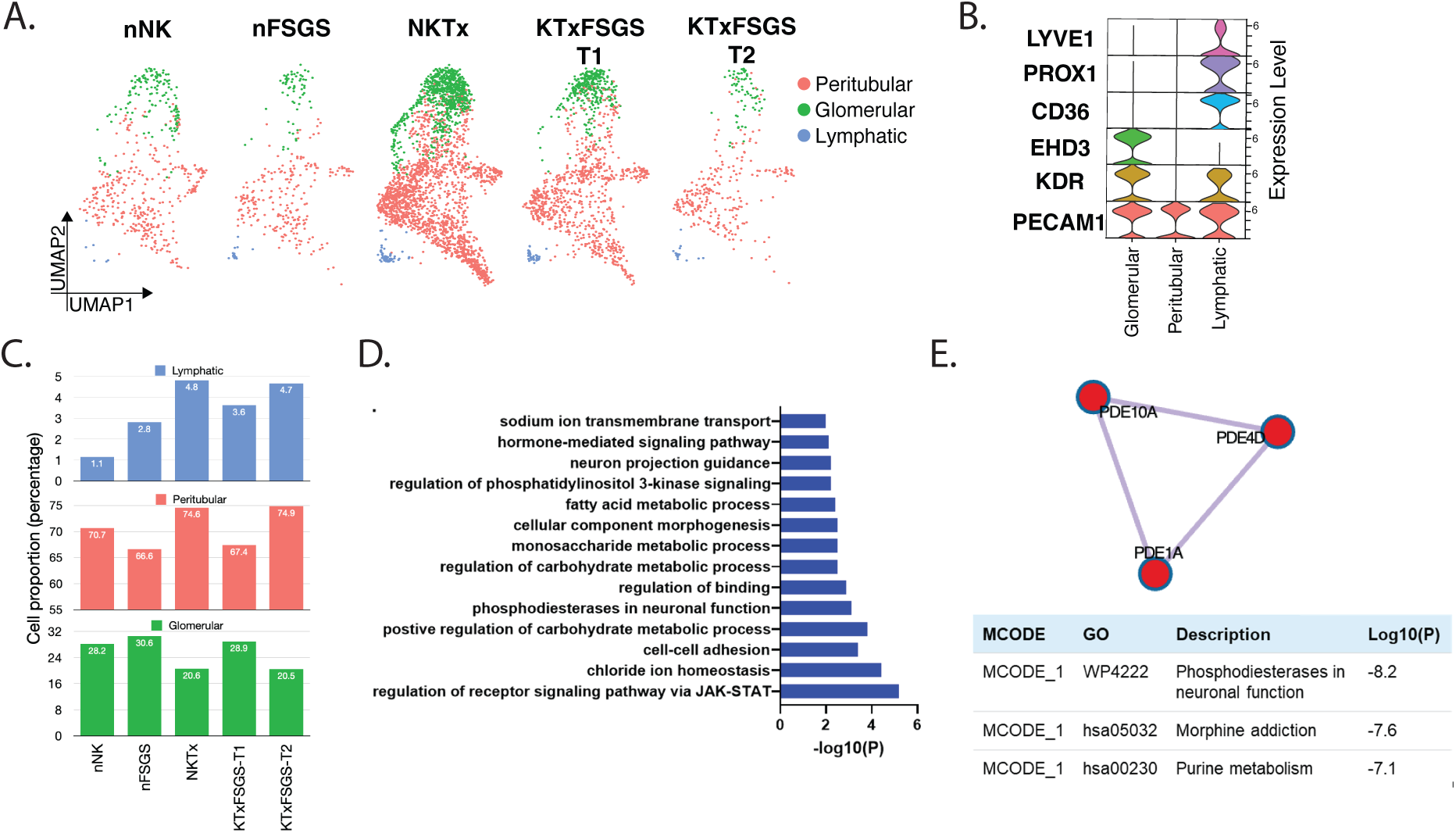
Pathway dysregulation in FSGS endothelial cells. (A) Endothelial cell clusters. (B) Gene markers in EC subclusters. (C) Proportion of ECs per condition. (D) Association of downregulated genes in TxFSGS-T1 EC with signaling pathways. (E) STRING interaction between phosphodiesterase in TxFSGS-T1.

### Proximal tubule cell injury, cell-type specific profiles, and fibrogenesis in posttransplant recurrent FSGS

The proximal tubule cluster cells were abundant, with 17,906 nuclei across two subclusters (PT-1 and PT-2, **Table S3**). The top gene markers for PT-1 were associated with solute carrier (SLC)-mediated transmembrane transport, metabolism, and other canonical functions of PT cells (**Fig. 5A**). In contract, the top gene markers for PT-2 were linked to plasma membrane-bounded cell projection assembly and actin filament organization (**Fig. 5B**). PT-2 cells expressed injury markers like *VIM*, *HAVCR1*, and *VCAM1*, indicative of tubule injury. These cells, described as adaptive PT (aPT), correlate with failed repair after kidney injury in both mice *(60)* and humans *(21)*. Notably, PT-2 cells also expressed *PROM1* and *CD24* (**Fig. 5C**), potentially indicating PEC differentiation to tubular progenitors, with bi-lineage potential demonstrated *ex vivo* and in mouse models *(46, 61, 62)*. The co-expression of *PROM1*, *CD24*, *VIM*, *HAVCR1*, and *VCAM1* in a subset of PT-2 cells suggests association with kidney regeneration and repair (**Fig. 5D**). In the PT-2 cell cluster of the NKTx group, 18.8% of cells co-expressed a subset of genes associated with both kidney injury and tubule-specific injury markers. In contrast, the KTxFSGS group exhibited 38% of such cells. This indicates that the KTxFSGS group has a larger subset of cells with failed repair potential.

**Fig. 5.**
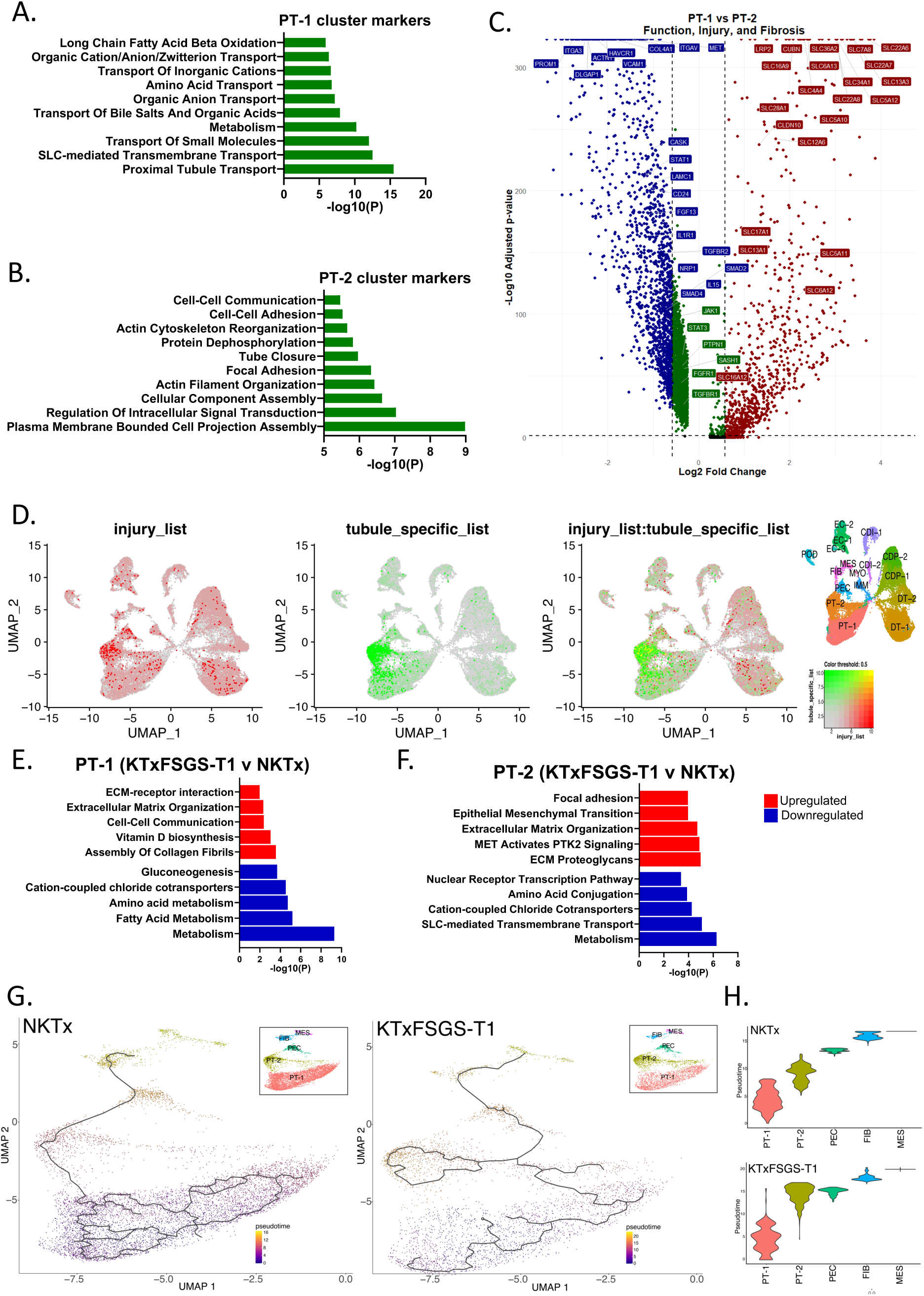
Proximal tubular cell injury, dysfunction, and fibrogenesis in FSGS. Gene Ontology of cellular markers characterizing (A) PT-1, and (B) PT-2. (C) Expression profile of genes associated with tubular injury, progenitor function, and fibrosis of PT-1 and PT-2 represented using Dot Plot. (D) Feature Plot to identify cells co-expressing (highlighted in yellow) both the kidney injury/progenitor markers (*PROM1, CD24, VIM*) and the tubule-specific injury markers (*HAVCR1, VCAM1*). (E) Gene Ontology and pathway analysis of DEGs between NKTx and KTxFSGS for (E) PT-1, and (F) PT-2. (G) Cell trajectory analysis of proximal tubule cell states in NKTx demonstrated homogeneity of cell states within the PT-1 cell population and overlap with the PT-2 cell population, while that of KTxFSGS-T1 showed overlapping cell states between injury PT-2 and PEC cell populations. Branched trajectories represent progression along different paths but isn’t necessarily related to real-life time. Pseudotime can be likened to a measure of progression along a continuum, such as stress states, where cells move in various directions over time. (H) This is also indicated by the highly overlapping pseudotime transcriptional profile between PT-2 and PEC in KTxFSGS-T1, indicating a similar stress state.

Differences in PT cell distribution between nFSGS and nNK were assessed as a baseline for evaluating KTxFSGS-T1. PT1 cells minimally changed in nFSGS (25.26% *vs.* 26.43% [nNK]) while PT2 increased significantly in nFSGS (16.74% vs. 3.56% [nNK]) (**Fig. 1C**). In KTx samples, the decrease in PT1 cells was more pronounced in KTxFSGS-T1 (18.15%) compared to NKTx (23.26%). Conversely, the proportion of PT2 cells increased in KTxFSGS (11.46%) compared to NKTx (3.72%). This indicates a decrease in functional PT cells and an increase in injured PT cells in KTxFSGS. In KTxFSGS-PT1 cluster, the upregulated DEGs showed an increased expression of collagens (*COL18A1*, *COL27A1*, *DST*, *COL4A1*) compared to NKTx, and an enrichment in genes involved in cell-cell communication and ECM organization. The downregulated DEGs in KTxFSGS PT1 cells enriched for pathways of metabolism, particularly fatty acid metabolism and peroxisomal lipid metabolism (**Fig. 5E**). The PT2 cluster associated DEGs showed an upregulation of extracellular matrix organization together with increased expression of COL23A1 and ECM proteoglycans (*ITGA3*, *ITGB1*, *ITGB3*, *ITGB5*, *ITGB6*, *LAMC1*, *LAMA5*, *VCAN*), potentially increasing cell adhesion and disrupting morphology of and around the tubular cells. In addition, a similar downregulation of metabolism-associated genes observed in the PT1 cluster was observed in PT2. This was accompanied by a downregulation of a myriad of SLC-mediated transmembrane transporters required for normal tubular function amino acid metabolism, and PPAR signaling pathway; all of which are required for normal kidney tubular cell function (**Fig. 5F**). In our trajectory analysis of cell state progression, we investigated the differentiation pathways and heterogeneity within PT-1 cell populations under varying conditions. This analysis revealed distinct branching patterns and stress responses in NKTx and KTxFSGS-T1 samples. In the NKTx samples, there was a demonstrated homogeneity of cell states within the PT-1 cell population, evidenced by multiple branching within PT-1 (**Fig. 5G**). However, in the KTxFSGS-T1 samples, a clearer branching out of trajectories from PT-1s to PECs was observed. This implies a distinct differentiation pathway or response to specific stimuli in the KTxFSGS-T1 condition. The pronounced branching suggests that disease-specific factors influence the transition from PT-1 to PEC, highlighting the heterogeneity in cell state progression and the impact of pathological conditions on cell fate determination. Furthermore, the presence of a highly overlapping pseudotime profile based on the transcriptome signature in PT cells between PT-2 and PEC demonstrated a similar “stress” state of these clusters (**Fig. 5H**). This stress state further emphasizes the impact of disease conditions on cell differentiation and behavior.

This led us to evaluate the differences between NKTx and individual KTxFSGS samples to identify time-associated injury differences in PT cells. In our dataset, the PT-2 cluster in all FSGS samples exhibited a significant increase in the expression of injury markers (*HAVCR1* and *VCAM1*, *VIM*) as well as progenitor-associated markers (*PROM1*, *CD24*; **Fig. S9**). Additionally, the PT1 cell in KTxFSGS-T1 displayed heightened expression of *HAVCR1*, and *PROM1*. Notably, in the sample with the shortest time post-KTx and first recurrence episode, FSGS2, there was an increase in the injury markers. However, in the T2 sample from the same patient there was an upregulation in *VIM* and *CD44* compared to other KTxFSGS samples. Enrichment pathway analyses of DEGs in PT1 of the FSGS2 patient showed upregulated genes that are associated with cell and tube morphogenesis, including collagens (*COL27A1, COL18A1*) and integrins (*ITGB3*, *ITGB6*, *ITGB8*, **Table S9, Fig. S9A**). However, all patients demonstrated a reduction in genes associated with the normal function of tubules in the PT-1. Interestingly, in all three patients, the downregulation of transport-associated functions was observed, with alterations in cell-cell adhesion and cell-junction organization. This indicates a loss of normal interactions and an increase in aberrant interactions in all the patients, regardless of the time or occurrence of the disease (**Fig. S9B and C)**. These findings support early tubular injury in PT cells in post-transplant FSGS recurrence. Critically, evidence of impaired repair and progression to fibrogenesis was also observed early in the disease recurrence and exacerbated with time and recurrence episodes.

### Increased B and T cell activation and distinct cDC pathways in KTxFSGS contribute to kidney injury and fibrosis

Sub-clustering of the immune cell cluster allowed the identification of multiple subclusters (**Fig. 6A and B**). nNK was excluded from the downstream analyses because of the low number of immune cells identified. A trend towards an increased proportion of CD4^+^ effector memory T cells (CD4_TEM_; cluster: CD4_TEM) (*ILR7*^+^ *IKT*^+^ *BCL11B*^+^ *TRAC*^+^) and B cells was observed in the KTxFSGS relative to NKTx. However, the trend in the increase of T cells was more pronounced in the follow up KTxFSGS-T2 (**Fig. 6C**). CD4_TEM_ and B cell populations increase in KTxFSGS-T1.

**Fig. 6:**
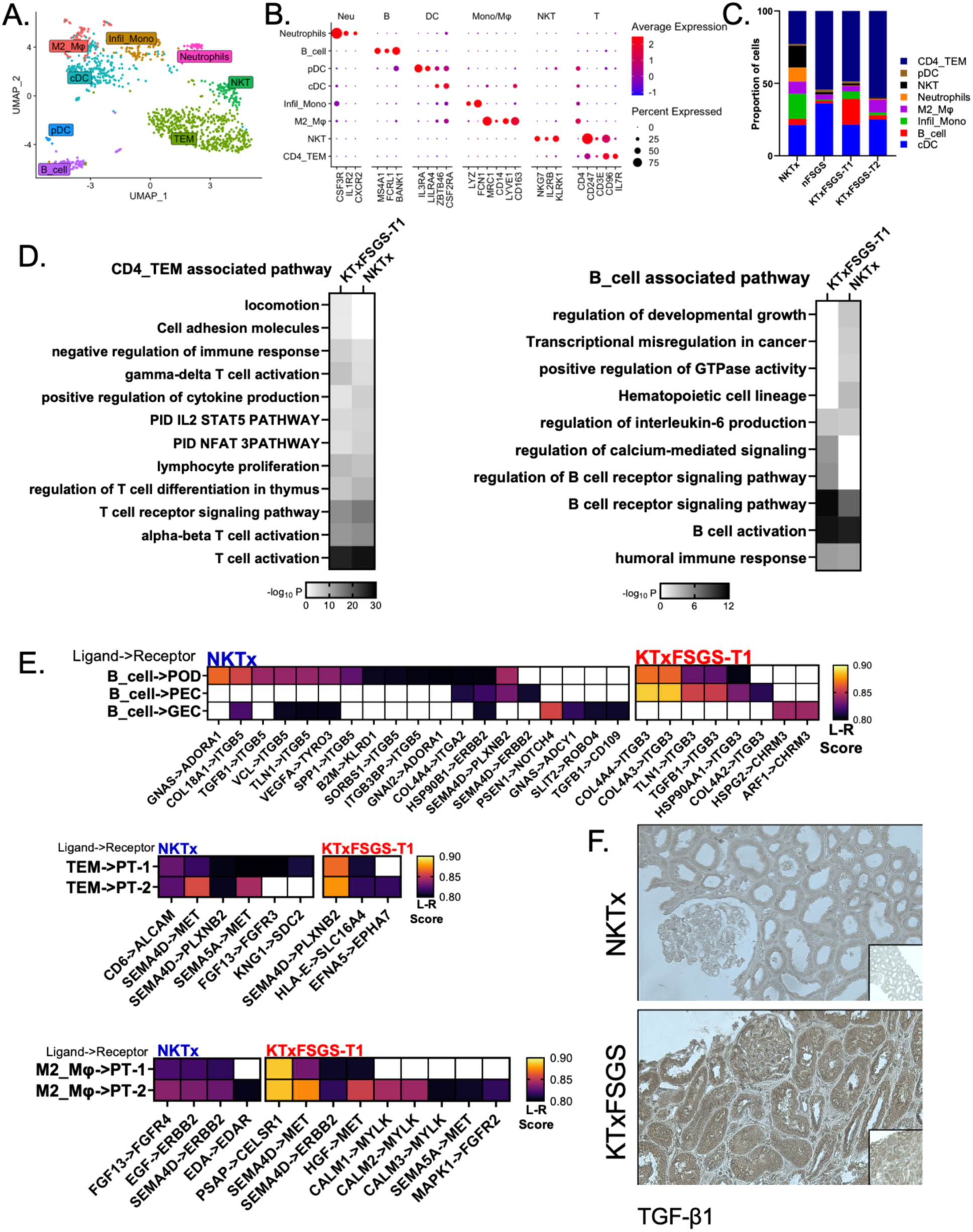
Molecular pathways associated with immune cells and their interaction with kidney cells. (A and B) UMAP of the immune cell clusters identified and marker genes used for the identification of the immune cell types in the dataset using the NKTx gene expression profile. (C) Relative proportion of immune cell subsets identified in each condition. (D) Major pathways associated with gene expression determined in two major immune cell types increasing in KTxFSGS-T1, CD4TEM and B cells, and (E) the major ligand-receptor interactions between the immune cells in NKTx and KTxFSGS-T1 with the cells of interest. (F) Immunohistochemical analysis of TGF-β1 in kidney biopsy from NKTx and KTxFSGS. The larger panel is at 20X magnification of a region of interest from within the smaller panel at 10X magnification.

*(63)*. GO and pathway analyses of the CD4_TEM_ cell gene marker signature identified enrichment of gamma-delta T cell activation, lymphocyte proliferation, and leukocyte aggregation in KTxFSGS-T1. The observed increase in gamma-delta T cells in FSGS is supported by a previous report, which also identified a significant presence of gamma-delta T cells in the native kidney with FSGS *(64)*. In contrast, in NKTx, there was an enrichment of alpha-beta T cell signaling pathways, positive regulation of cytokine production, regulation of RUNX1 expression and activity, and MAPK family signaling cascades (**Fig. 6C**). Both conditions showed an increase in T cell receptor signaling pathway and T cell activation. This suggests that while both conditions have an active T cell population, inherent differences in cellular properties exist, with increased proinflammatory signaling during KTxFSGS. Interestingly, the NKTx profile indicates a proinflammatory signature that has been described to persist in normally functioning long-term allografts *(21)*. Similar increased activation was observed in the B cell cluster for KTxFSGS-T1. Gene markers of B cells enriched for humoral immune response, B cell activation, and regulation of B cell receptor signaling (**Fig. 6D**). The potential increase in activation and regulation of signaling in KTxFSGS-T1 samples is highly relevant due to the observations demonstrating a role of B cells in the pathogenesis of FSGS *(65–67)*. NKTx presented an overall higher proportion of total macrophages/monocytes, further identified as <u>M2</u>-like <u>Mφ</u> (Cluster: M2_Mφ), and <u>infil</u>trating <u>mono</u>cytes (Cluster: Infil_mono). It was interesting to note that the M2_Mφ is a tissue-resident macrophage, positive for *LYVE1*, *MRC1*, *CD163*, and characterized as “M2-like” due to the expression of *MSR1*, *TGFBR1*, and *CD200R1*. While the role of M2 macrophages is not fully established and they may be involved in chronic rejection, a beneficial reparative role of M2 macrophages in kidney allografts has also been reported *(68)*. Conventional DC (cDC) were another immune cell cluster of interest in our dataset, as they were the most prominent innate immune cell cluster in the KTxFSGS-T1 samples. A marker gene-based over representation analysis (ORA) demonstrated enrichment of genes associated with regulation of cell activation, endocytosis, response to cytokine stimulus, inflammatory response, and neutrophil degranulation in the KTxFSGS-T1 samples. A negative role of DCs in the pathogenesis of kidney injury and disease has been documented *(69)*. The depletion of DCs was observed to impair the activation and proliferation of T cells in kidney injury *(70, 71)*. Moreover, an increased presence of DCs in kidney biopsies of patients with FSGS has been observed *(71)*.

Next, we determined the interaction between the immune cells and cells of the glomerulus and tubular cells in NKTx and KTxFSGS-T1, utilizing the RNA expression and an *in-silico* database. As expected, there were multiple overlapping interactions between immune cells and kidney cells in both NKTx and KTxFSGS-T1; however, some interactions were unique to KTxFSGS-T1 and could potentially explain the reported increase in B and T cells during recurrent FSGS. For example, multiple B cell-induced ligands, including type 4 collagens, and *TGFB1* could interact with *ITGB3* on both podocytes and PECs. This is particular important because activation of *ITGB3* signaling in multiple studies has been demonstrated to induce senescence and fibrosis in cells, including tubular cells *(72–74)*. Indeed, an increased TGFβ1 immunostaining in surviving glomeruli of diseased kidneys of KTxFSGS relative to NKTx was also observed (**Fig. 6F**). Furthermore, in NKTx, T cells could induce MET signaling in the PT cells, which is known to decrease in transplant glomerulopathies and acute kidney injury *(75, 76)*. Similarly, the M2_Mφ could interact with the PT cells to increase ERBB2 signaling, whereas in KTxFSGS-T1, they could promote EMT and fibrosis through activation of MYLK and FGFR2 signaling.

## Discussion

FSGS recurrence is a significant concern in kidney transplant recipients. Despite advancements in immunosuppressive therapies, FSGS recurs in 25-50% of first transplants and up to 80% in subsequent transplants *(77)*. Previous studies have suggested that a circulating permeability factor plays a key role in FSGS recurrence, but the exact mechanisms remain unclear *(78, 79)*. Clinicians manage recurrent FSGS through plasma exchange, immunosuppressive therapies, and supportive care measures *(80)*. However, a deeper understanding of the molecular and cellular mechanisms underlying FSGS recurrence is essential for developing more effective treatments *(81)*.

The unique features of our study include (i) a distinct patient population tested, specifically KTRs with post-transplant recurrent idiopathic FSGS, with follow-up samples from two patients during subsequent recurrences; (ii) identification of molecular alterations within individual kidney cell type clusters; (iii) a high proportion of parenchymal cell recovery; and (iv) identification of potential renal-immune cell and cell-matrix interactions. These findings provide critical insights that could inform future research and therapeutic strategies.

Podocytes primarily rely on glycolysis for energy and lipid metabolism for the development of secondary metabolites and signaling molecules, essential for maintaining the active cytoskeleton (**Fig. 7**) *(82–85)*. The downregulation of *XYLB*, crucial for the pentose phosphate pathway from xylulose, suggests a disruption in glucose and lipid metabolism, which could lead to podocyte dysfunction and detachment *(86–88)*. Indeed, there was a downregulation of multiple genes associated with lipolysis, such as *PTGER4*, *PTGER3*, and *PTGDS*. *PTGER3* has been demonstrated as one of the major hub proteins inhibiting diabetic nephropathy-induced podocyte apoptosis downstream the activated PPARγ signaling pathway *(89)*. In fact, *PTGDS* was observed to be significantly downregulated following injury in podocytes representing IgA nephropathy *(90)*. Previous studies have identified loss-of-function mutations in *PLCE1* as contributing factors to genetic FSGS in pediatric patients, underscoring its critical role in recurrent FSGS*(91)*. Metabolic changes have also been observed in the podocytes from other glomerular conditions during diseased condition. Interestingly, the gene expression profile differences have been relatively different between conditions suggesting towards a disease dependent podocyte gene expression modulation. For example, a glomerulosclerosis model in mice, after nephrotoxic immune injury, podocytes showed acute activation of the Hippo pathway with an increased gene expression profile for oxidative phosphorylation and triglyceride homeostasis *(92)*. In fact, it has been demonstrated there is a highlighted glucose-independent responses in glomerular cells and mechanosensitive signaling pathways during diabetes-induced nephropathies *(93)*. During chronic kidney disease in obesity-related glomerulopathy, there is an enrichment in genes related to oxidative phosphorylation, cell adhesion, thermogenesis, inflammatory pathways, and fluid shear stress in obesity-related glomerulopathy *(63)*. The prevalence of downregulated genes points to a general decline in certain cellular functions within podocytes, likely reflecting a state of cellular stress or injury. Reduced expression of *XYLB*, encoding xylulokinase (XK), leads to loss of the “xylulose-5-phosphate (X5P) formation” pathway, critical for glycolysis and glucose-induced lipogenesis—key energy pathways for podocytes *(82, 83, 86, 87)*. Additionally, decreased expression of *PPARGC1A* results in reduced activation of PGC-1alpha, which is vital for maintaining podocyte homeostasis *(84)*. Previous reports have shown reduced expression of PPARGC1A in both diabetic and non-diabetic chronic kidney disease *(94)*.

**Fig. 7:**
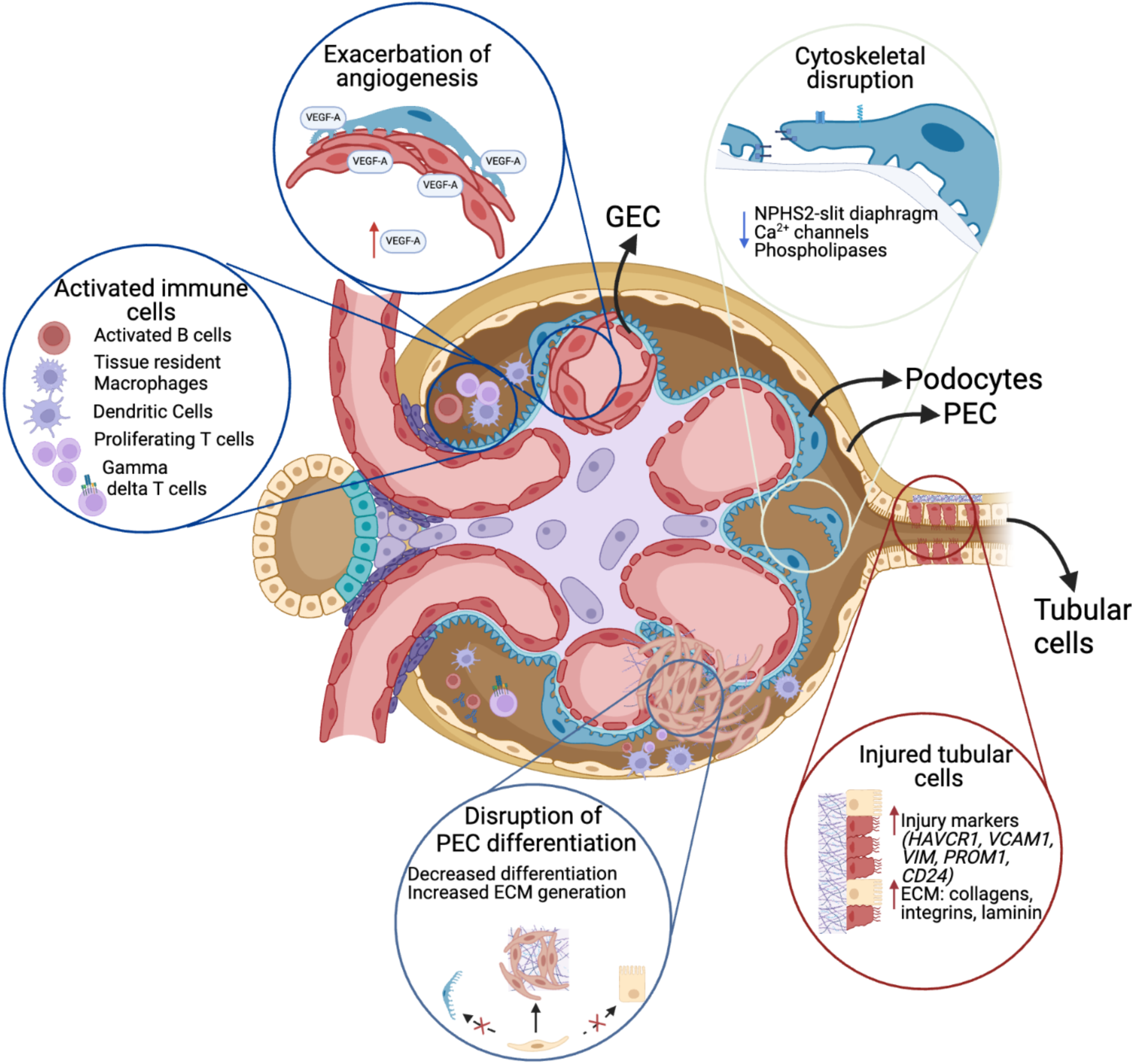
Molecular and Cellular Alterations in KTxFSGS. The recurrence of FSGS is driven by a complex cascade of injuries and changes within the glomerulus. These include an increase in glomerular endothelial cells (GECs), reduced differentiation of parietal epithelial cells (PECs) into podocytes or tubular cells, and elevated extracellular matrix (ECM) production. These changes, along with immune cell infiltration, contribute to glomerular sclerosis. Concurrently, podocytes experience cytoskeletal disruption, leading to their detachment and resulting in proteinuria. The elevated proteinuria, in turn, damages proximal tubular cells (shown in red), triggering an injury response that exacerbates tubular sclerosis and ultimately leads to graft loss. Created using BioRender.

The cooperative and individual roles of calcium ion transporters and phospholipases are crucial for maintaining normal podocyte function. When podocytes are subjected to inflammatory stimuli, the activation of PLC cascades can lead to an increased intracellular Ca^2+^ concentration, which further activates pathways that regulate cytoskeletal dynamics and cell survival *(95)*. Recent reports have demonstrated that mutation in *PLCE1* causes recessive nephrotic syndrome *(96)*. Disruption of calcium homeostasis can lead to podocyte detachment by affecting cytoskeletal dynamics, impairing adhesion molecules, compromising cell-cell junctions, triggering apoptosis, and increasing reactive oxygen species (ROS) production *(95, 97–101)*.

Disruption of podocyte foot processes is intimately linked to perturbations in calcium homeostasis, which, alongside alterations in prostaglandin and inositol 3-phosphate signaling, serve as crucial secondary messengers in podocyte function. In our study, we observed that these disruptions were exacerbated by the downregulation of calcium channels, *LRPC1* and *LRPM1*, during KTxFSGS, highlighting a potential mechanistic pathway contributing to podocyte injury and subsequent disease progression. Interestingly, our findings revealed an upregulation of *PODXL* and *NPHS1* in KTxFSGS podocytes. While the increase in *NPHS1* likely represents a compensatory response to heightened injury and shear stress, the aberrant upregulation of *PODXL* is particularly noteworthy. This ectopic expression of *PODXL* has been documented in various pathological states and is associated with compromised podocyte integrity *(102, 103)*. The antiadhesive properties of PODXL protein are responsible for keeping the intercellular spaces open *(104, 105)*, further contributing to podocyte dysfunction, potential detachment, and disease progression. This aligns with the work reported by Latt et al. *(20)*, who reported the presence of viable podocytes exhibiting an EMT signature.

Notably, the urine cell pellet of patients with FSGS contained a significant proportion of macrophages *(20)*, further supporting our observation of reduced macrophage proportion in KTxFSGS tissues. The upregulation of CD44 in podocytes only during early pathological changes and subsequent metabolic disturbances likely contribute to podocyte depletion and proteinuria, eventually leading to renal failure. Moreover, while it has been hypothesized that the podocyte loss could be a result of detachment from the GBM, cell death could also be a cause of reduction in POD. Despite being a cross-sectional study, the gene expression profiles did not show an enrichment of genes responsible for cell death related pathways in FSGS.

Podocytes are terminally differentiated epithelial cells, and their replenishment has been demonstrated to occur through the dedifferentiation of glomerular PECs *(106, 107)* CD44, a transmembrane glycoprotein, plays a role in cell interactions, adhesion, and migration. It is typically absent in healthy podocytes but upregulated in response to glomerular injury and inflammation. CD44 was identified only in the earlier samples with FSGS recurrence and was absent in later samples. Our data indicates that during the early phase of recurrent FSGS, there is a significant increase in CD44 expression. However, CD44 activation was not observed in NKTx, nNK, or transplanted kidneys without recurrent FSGS. This suggests that the initial podocyte response in recurrent FSGS is to increase CD44 expression, a response that diminishes in later episodes, highlighting the temporal progression of the disease. Our analysis suggests that while podocyte loss is a hallmark of FSGS, significant alterations in genes associated with cell cycle arrest or apoptosis are not discernible. Instead, we observed changes in genes essential for metabolic homeostasis and structural integrity. This indicates that podocyte depletion in FSGS recurrence post-transplant predominantly arises from metabolic dysregulation, particularly related to lipid metabolism and structural compromise.

The presence of regenerative PECs, glomerular scarring was observed, with increased levels of ECM collagens in PECs during recurrent episodes of FSGS. Marker genes associated with regeneration were reduced as recurrence episodes increased. PECs also showed increased ECM deposition and fibrosis, aligning with previous studies that highlight their dual role in podocyte regeneration and disease progression *(108)*. The increased VEGF-A release from PODs in response to injury and the subsequent response of ECs to reduce permeability further contribute to the glomerular injury and restructuring of the microvasculature. This is a unique observation for reFSGS in our patient population. In a previous pre-clinical study using mutant Wt1, the early stages of Wt1^R394W/+^ disease showed disrupted angiogenic signaling between podocytes and glomerular endothelial cells. This was observed due to a reduction in the expression of *Vegfa* and *Nrp1*, in the podocytes resulting in the loss of endothelial cells and the development of WT1 glomerulopathies *(109)*.

Interestingly, in the reFSGS samples, ECs exhibited increase in glomerular EC, correlating with vascular changes in FSGS pathology. The study identified three unique clusters of ECs, with changes in glomerular and peritubular endothelial cells. Downregulated genes associated with cellular mobility and vascular integrity were noted, indicating endothelial dysfunction *(110)*. This supports findings that link endothelial injury to proteinuria and glomerular sclerosis, emphasizing the role of endothelial cells in FSGS recurrence *(18)*. Glomerular ECs were also found to contribute to the development of IgA nephropathy (IgAN) in a preclinical mouse model of IgAN using scRNAseq. The authors observed and enrichment of “antigen processing and presentation of peptide antigen via MHC class Ib,” and “T-cell migration” in the IgAN glomerular ECs *(111)*. Similarly, another scRNA-seq study on IgAN identified the potential activation of podocytes and glomerular ECs through the slit receptors *(111)*. Again, these was in contrast to our studies where we observed reduction in metabolic activities, STAT activation, and reduced cell-cell adhesion. PTs showed injury markers and increased ECM gene expression, indicating early tubular injury and impaired repair mechanisms in FSGS recurrence. Increased injury markers and progenitor markers in PT cells were observed early in the disease progression, with these cells contributing to ECM production and fibrosis activation. These findings underscore the importance of PT cells in chronic kidney disease progression and align with previous research on tubular cell stress responses.

We acknowledge the limitations of our study, particularly the small number of biopsies. However, the use of snRNAseq provides a powerful approach for generating high-dimensional data from a large number of individual cells. This method enabled us to analyze over 50,000 individual cells across the biopsy samples, providing a high-resolution view of the cellular landscape. This large cell number significantly enhances the sensitivity of our analysis, allowing us to identify subtle molecular and cellular processes involved in FSGS recurrence, even within a small sample size. The extensive dataset enables a more granular exploration of cell-type-specific changes and disease mechanisms that would be difficult to capture with traditional methods. The large-scale cellular data allows for a comprehensive exploration of cellular heterogeneity and the mechanisms underlying disease recurrence. Thus, the single-cell data provides robust insights into gene expression patterns, cell-type identification, and pathway activity across many cells. Overall, the use of this technology enhances the statistical power of the observations and provide a high resolution to capture rare cell populations, identify subtle gene expression differences, and detect cell-type specific changes that would not be evident in a smaller sample with traditional methods.

While our study did not directly assess the presence of circulating auto-antibodies, the canonical slit diaphragm genes such as *NPHS1* (nephrin) had a slight increase in expression as opposed to previous studies that reported a reduction in NPHS1 in presence of NPHS1 auto-antibodies *(112)*. This may reflect sample-specific variability or in this case may suggest that other mechanisms beyond auto-antibody-mediated injury contributing to the observed molecular changes. Given the established link between auto-antibodies and podocyte damage, the absence of detectable regulation of slit diaphragm genes in our analysis warrants consideration as a study limitation. However, it is to be noted that at present there is insufficient evidence to support the role of auto-antibodies as the circulating “factor” that triggers FSGS and their use as a prognostic biomarker for reFSGS *(113)*. Similarly, the non-significant difference in suPAR expression suggests that these patients may not have it as a circulating factor causing the recurrence of disease.

This first study evaluating posttransplant FSGS recurrence at the single cell level, provides novel insights into the cellular and molecular mechanisms of FSGS recurrence post-KT. The identification of transcriptional changes in podocytes, PECs, ECs, and tubular cells highlights the complexity of FSGS and the potential role of circulating factors. The findings underscore the importance of targeted therapies to address specific cellular dysfunctions and improve outcomes for patients with recurrent FSGS. By advancing our understanding of FSGS at the single-cell level, this research lays the groundwork for future studies aimed at developing precision medicine approaches for kidney disease. This study provides the evidence for performing clinical studies with longitudinal biopsies and peripheral blood collections from KTx recipients ranging from pre-transplant to recurrence of disease to identify the cellular and molecular profiles during the development of reFSGS.

## MATERIALS AND METHODS

### Patients and samples

Clinical and demographic information of the patient population used in the study has been outlined in **Tables 1 and 2**. All patients received triple-drug immunosuppression that included tacrolimus, mycophenolate mofetil, and prednisone. As control groups, we included three normal native kidneys (nNK) (GSE131882 *(23)*) and four kidney grafts with normal/stable graft function (NKTx) whose biopsies were categorized as normal/non-specific. NKTx patients were recruited as part of a study that included surveillance biopsies. The Institutional Review Board approved the study and patients signed informed consent at the time of transplantation (HP-00091954). Patients with KTxFSGS had biopsies done at the time of FSGS recurrence. NKTx were >15 months post-transplantation, had an estimated GFR (eGFR) of ≥60 mL/min/1.73m2, no proteinuria, no circulating IgG antibodies against donor HLA at the time of biopsy, and had normal/non-specific findings in the allograft surveillance biopsies.

Recurrent focal segmental glomerulosclerosis (FSGS) was diagnosed in the transplant recipients based on clinical, histopathological, and clinical history findings. These patients had a known history of FSGS as the underlying cause of end-stage renal disease (ESRD) in their native kidneys. Clinically, all patients exhibited significant nephrotic-range proteinuria (≥3.8 g/day) and impaired renal function, with elevated serum creatinine and reduced eGFR. Histopathological examination revealed typical features of FSGS recurrence, including diffuse effacement of podocyte foot processes, cytoskeletal changes, and microvillous transformation in severe cases, without evidence of immune complex deposition (negative C4d). However, one patient showed anti-C4d positivity at the time of follow-up recurrence, indicating a possible antibody-mediated rejection (AMR) in addition to recurrent FSGS. The temporal association of these findings with prior transplant timelines and recurrent episodes further supported the diagnosis of recurrent FSGS, though the presence of anti-C4d necessitates close monitoring for potential rejection. Kidney biopsy tissue samples were preserved in RNA*later* at the site of biopsy for sequencing analysis and in ethanol for downstream processing for immunohistochemical analysis. The clinical and research activities being reported are consistent with the Principles of the Declaration of Istanbul as outlined in the ‘Declaration of Istanbul on Organ Trafficking and Transplant Tourism’.

### Sample processing and transcriptomic sequencing

Nuclei isolation was conducted as per our established protocol *(107)*. Isolated nuclei were processed using Chromium Single Cell 5’ Reagent Kits as per the manufacturer’s protocol. RNA was reverse transcribed into cDNA within individual droplets. cDNA concentration was quantified using a Qubit fluorometer (ThermoFisher), with approximately 20ng total added to each reaction. After breaking the emulsion, cDNAs were amplified and fragmented, followed by the addition of Illumina adapters using the 10x Chromium Single Cell 5′ Library & Gel Bead Kit (v2) according to the manufacturer’s instructions (10x Genomics). The reactions were conducted at 98°C for 45 seconds, followed by 16 cycles of 98°C for 20 seconds, 67°C for 30 seconds, and 72°C for 1 minute, with a final extension step at 72°C for 1 minute, and then held at 4°C. Post-PCR, cDNA quantity was determined again using a Qubit fluorometer (ThermoFisher). Analysis with a Bioanalyzer (Agilent) and gel electrophoresis were employed to confirm the expected size post-adapter ligation. Finally, the multiplexed libraries were combined and sequenced on the S4 flow cell of a NovaSeq 6000 (Illumina) using 150bp paired-end sequencing.

### snRNA-seq data analysis

The FastQ files from the 10x Genomics protocol were aligned to the human pre-mRNA reference sequence (GRCh38) using CellRanger (v3). Quality control (QC) assessed the number of detected features, gene expression depth, and mitochondrial gene expression. Genes expressed in fewer than three cells, cells with fewer than 400 or over 5,000 expressed genes, and nuclei with over 2.5% mitochondrial gene expression were excluded. Samples were integrated using ‘Seurat’ and tested for batch effects. Cell clustering was done using PCA and UMAP. Distinct cell clusters and cluster-specific gene markers were identified. These markers were compared to a database to assign potential cell types. Major and unknown cell types were identified. Cell type proportions were analyzed for population differences among conditions. Further analyses included subclustering functionally relevant cell types, assessing differentially expressed genes (DEGs), and identifying enriched gene ontology terms and functional pathways (FDR ≤0.05 and FC ≥1.5). All analyses and visualizations were conducted using R.

### Gene ontology analysis

Enrichr and Metascape were used for over representation analysis to compare intra- and inter-cell clusters between the study groups. Pathways with significant enrichment were identified using an p value threshold of ≤0.05. Differentially expressed genes (DEGs) were further analyzed with WebGestalt23, SHINYGO or DAVID (v6.8) to map genes to relevant biological functions.

### Single-cell trajectory analysis

Single-cell dynamics were investigated using Monocle v3.30, an unsupervised algorithm that orders single-cell expression profiles in pseudotime, allowing for multiple cell fates from a single progenitor cell type. For each condition, the Seurat SummarizedExperiment object was converted into a CellDataSet object for Monocle3. The gene expression matrix identified cell clusters, which were grouped into cell partitions. Trajectory graphs for predetermined cell types were inferred based on gene expression changes, positioning cells along a pseudo timeline as they transitioned between states. Monocle also modeled changes in gene expression over pseudotime. Significant pseudotime-related genes were selected based on an adjusted p-value < 0.05, Moran’s I statistic > 0, and expression in at least 10% of cells. Visualizations were created using R.

### Ligand receptor analysis

To investigate potential cell-cell interactions among immune cells and kidney cell types across the conditions we used ligand-receptor pair data from LRdb2. The interaction score (LRscore) was calculated using an expression product model that incorporates expression values for interacting proteins from two distinct cell types, regularized with the mean expression matrix to yield a pseudo-likelihood ratio between 0 and 1. We applied a score threshold of 0.5 and retained LR pairs where either the ligand or receptor was expressed in at least 25% of the cells. Visualizations were created using GraphPad Prism.

### Immunohistochemistry

Formalin-fixed paraffin-embedded (FFPE) kidney tissue sections from NKTx and KTxFSGS patients were analyzed to assess the expression of EGF (MA5-15571) and TGFβ1 (MA5-16949) via DAB-based chromogenic immunohistochemistry (IHC). Standard IHC protocols were followed, including antigen retrieval using acidic citrate buffer under pressurized heat, followed by endogenous peroxidase quenching with 3% hydrogen peroxide. For co-localization studies, fluorescence-based IHC was performed to evaluate the co-expression of PODXL and CLDN1. Antigen retrieval and antibody labeling were carried out using standard protocols in accordance with manufacturer guidelines (Novus Biologics).

## Data availability

The single cell RNA sequencing data of the kidney biopsies generated in this study have been deposited in the GSE269078.

## List of Supplementary Materials

### Supplementary Figures

**Fig. S1: Quality control parameters of the study.** These included (A) percent mitochondrial gene expression per cell (percent.mito) versus the total number of RNA detected per cell (nCount_RNA), (B) the number of genes detected in cell (nFeature_RNA) versus nCount_RNA, also depicted are (C) nFeature_RNA, nCount_RNA, and percent.mito per sample. The cut off used in the study is present in the main text.

**Fig. S2: Cell cluster identification.** (A) Total cell clusters identified in our single nuclei (sn) RNA sequencing presented as an integrated uniform manifold approximation and projection (UMAP). (B) The integrated UMAP was split based on patient condition consisting of normal native kidney - nNK, kidney allograft with normal function - NKTx, native kidney with focal segmental glomerulosclerosis (FSGS) - nFSGS, transplanted kidney with recurrent FSGS at first collection – KTxFSGS-T1, and second collection during the next recurrence - KTxFSGS-T2. Nineteen clusters of cells including the two clusters that had markers of multiple cell types (UNK-1 and UNK-2) which were excluded from the study in our current downstream analysis.

**Fig. S3: Treatment of conditionally immortalized podocytes with serum of patients with recurrent FSGS.** Podocytes were grown under immortalization conditions as per the vendor’s instructions after which they were allowed to differentiate for 5 days in the differentiation media. On the 5^th^ day, the cells were cultured with media supplemented with 4% of patient serum (N = 2) and cultured for additional 48 hours after which the RNA was collected and Taqman based qPCR (n=2) was used to determine expression of genes mentioned in the figure, p-value * < 0.05 (Student’s t-test).

**Fig. S4: Biological properties associated with differentially expressed genes between NKTx and KTxFSGS.** (A) EnrichR based identification of pathways associated with significantly downregulated genes in podocytes of KTxFSGS-T1 compared to KTx. (B)DotPlot of cell death related genes in NKTx (Normal) and post-transplant recurrent FSGS (PostTXwFSGS) does not demonstrate an enrichment of apoptosis-associated genes in either condition.

**Fig. S5:** Metascape-based pathway analysis of differentially expressed genes in podocytes of individual recurrent T1-FSGS patient samples (FSGS1-3) compared to normal allograft (NKTx)

**Fig. S6:** DotPlot demonstrating the presence of CD44 in podocytes cluster (POD) of FSGS2 (POD_GEX-NW-5). Classical podocyte associated genes have also been plotted for reference.

**Fig. S7:** DotPlot demonstrating the presence of CD44, HB-EGF, and EGF in PEC cluster, represented in (A) all conditions, and (B) all normal allograft samples.

**Fig. S8:** Metascape-based pathway analysis of differentially expressed genes glomerular endothelial cells.

### Supplementary Tables

**Table S1:** Summary of Illumina sequencing results

**Table S2:** Summary of Q30 scores

**Table S3:** Counts of isolated cells in each cell cluster

**Table S4:** Top markers for cell clusters

**Table S5:** List of cluster-based gene expression markers used to determine cluster identity

**Table S6:** List of differentially expressed genes between KTxFSGS-T1 and NKTx

**Table S7:** List of differentially expressed genes between KTxFSGS-T1 and nFSGS

**Table S8:** List of differentially expressed genes between T1-FSGS1 and NKTx

**Table S9:** List of differentially expressed genes between T1-FSGS2 and nKTx

**Table S10:** List of differentially expressed genes between T1-FSGS3 and NKTx

**Table S11:** List of differentially expressed genes between nNK and nFSGS - Glomerular Endothelial Cell cluster only

**Table S12:** List of differentially expressed genes between NKTx and KTxFSGS-T1 - Glomerular Endothelial Cell cluster only

## Acknowledgements

We would also like to thank the members of the Institute for Genome Sciences at the University of Maryland, Baltimore for high-throughput sequencing support. We also appreciate the support of Kirk Campbell, MD at the Icahn School of Medicine at Mount Sanai for his help with podocyte cell cultures. We also like to thank former members of the Mas Lab, Jennifer McDaniels, PhD; Cem Kuscu, PhD; and Canan Kuscu, PhD for their support with initial set up of the single cell RNA sequencing assays.

## Funding

This research reported in this publication was supported by the

National Institutes of Health R21DK100678 (DGM)

National Institutes of Health R01DK109581 (VRM)

National Institutes of Health R01DK122682 (VRM)

National Institutes of Health R21AI172077 (VRM).

University of Maryland Baltimore, Institute for Clinical & Translational Research (ICTR)/Clinical and Translational Science Award (CTSA) grant ULTR003098 (HZ)

## Author contributions

Conceptualization: VRM, LG, DGM, HZ

Methodology: TVR, SA, HZ, ACS, EB

Investigation: VRM, LG, ACS, TVR, SA, DGM, HZ, CBD

Visualization: TVR, ACS, HZ, EB

Funding acquisition: VRM, DGM, LG

Project administration: VRM, DGM, LG

Supervision: VRM, LG, DGM

Writing – original draft: VRM, TVR, HZ, ACS

Writing – review & editing: VRM, TVR, HZ, ACS, SA, CBD, EA, MT, DGM, EF

## Data and materials availability

All data, code, and materials used in the analysis must be available in some form to any researcher for purposes of reproducing or extending the analysis. Include a note explaining any restrictions on materials, such as materials transfer agreements (MTAs). Note accession numbers to any data relating to the paper and deposited in a public database; include a brief description of the data set or model with the number. If all data are in the paper and supplementary materials, include the sentence “All data are available in the main text or the supplementary materials.”

## References

1. K. D. Guruswamy Sangameswaran, M. F. Hashmi, K. M. Baradhi, in StatPearls, (StatPearls Publishing, Treasure Island (FL), 2024).

2. M. Sambharia, P. Rastogi, C. P. Thomas, Monogenic focal segmental glomerulosclerosis: A conceptual framework for identification and management of a heterogeneous disease. Am J Med Genet C Semin Med Genet 190, 377–398 (2022).

3. M.-H. Han, Y.-J. Kim, Practical Application of Columbia Classification for Focal Segmental Glomerulosclerosis. Biomed Res Int 2016, 9375753 (2016).

4. J. L. Colquitt, J. Kirby, C. Green, K. Cooper, R. S. Trompeter, The clinical effectiveness and cost-effectiveness of treatments for children with idiopathic steroid-resistant nephrotic syndrome: a systematic review. Health Technol Assess 11, iii–iv, ix–xi, 1–93 (2007).

5. P. A. S. Vaz de Castro, A. A. Amaral, M. G. Almeida, H. Selvaskandan, J. Barratt, A. C. Simões E Silva, Examining the association between serum galactose-deficient IgA1 and primary IgA nephropathy: a systematic review and meta-analysis. J Nephrol (2024), doi:10.1007/s40620-023-01874-8.

6. Y. Zhuang, H. Lu, J. Li, Advances in the treatment of IgA nephropathy with biological agents. Chronic Dis Transl Med 10, 1–11 (2024).

7. Z. Chen, D. Xu, F. Cui, H. Hou, Z. Mao, X. Gao, Coexistence of anti-glomerular basement membrane disease and IgA nephropathy: an illustrative case and comprehensive literature review. Ren Fail 46, 2323160 (2024).

8. A. D. Kistler, D. J. Salant, Complement activation and effector pathways in membranous nephropathy. Kidney Int 105, 473–483 (2024).

9. C. Murtas, M. Bruschi, S. Spinelli, X. Kajana, E. E. Verrina, A. Angeletti, G. Caridi, G. Candiano, S. Feriozzi, M. Prunotto, G. M. Ghiggeri, Novel biomarkers and pathophysiology of membranous nephropathy: PLA2R and beyond. Clin Kidney J 17, sfad228 (2024).

10. L. J. Hickson, M. Gera, H. Amer, C. W. Iqbal, T. B. Moore, D. S. Milliner, F. G. Cosio, T. S. Larson, M. D. Stegall, M. B. Ishitani, J. M. Gloor, M. D. Griffin, Kidney Transplantation for Primary Focal Segmental Glomerulosclerosis: Outcomes and Response to Therapy for Recurrence. Transplantation 87, 1232 (2009).

11. M. Rudnicki, FSGS Recurrence in Adults after Renal Transplantation. BioMed Research International 2016, e3295618 (2016).

12. T. Törnroth, Recurrent and de novo glomerulonephritis in allografted kidneys: aspects of ultrastructural diagnosis. Appl Pathol 5, 88–94 (1987).

13. M. Ismail-Allouch, G. Burke, J. Nery, D. Roth, V. Esquenazi, P. Ruiz, J. Miller, Rapidly progressive focal segmental glomerulosclerosis occurring in a living related kidney transplant donor: case report and review of 21 cases of kidney transplants for primary FSGS. Transplant Proc 25, 2176–2177 (1993).

14. A. Uffing, M. J. Pérez-Sáez, M. Mazzali, R. C. Manfro, A. C. Bauer, F. de Sottomaior Drumond, M. M. O’Shaughnessy, X. S. Cheng, K.-K. Chin, C. G. Ventura, F. Agena, E. David-Neto, J. B. Mansur, G. M. Kirsztajn, H. Tedesco-Silva, G. M. V. Neto, C. Arias-Cabrales, A. Buxeda, M. Bugnazet, T. Jouve, P. Malvezzi, E. Akalin, O. Alani, N. Agrawal, G. La Manna, G. Comai, C. Bini, S. A. Muhsin, M. C. Riella, S. R. Hokazono, S. S. Farouk, M. Haverly, S. S. Mothi, S. P. Berger, P. Cravedi, L. V. Riella, Recurrence of FSGS after Kidney Transplantation in Adults. Clin J Am Soc Nephrol 15, 247–256 (2020).

15. M. A. Saleem, Molecular stratification of idiopathic nephrotic syndrome. Nat Rev Nephrol 15, 750–765 (2019).

16. J. B. Kopp, H.-J. Anders, K. Susztak, M. A. Podestà, G. Remuzzi, F. Hildebrandt, P. Romagnani, Podocytopathies. Nat Rev Dis Primers 6, 68 (2020).

17. S. Hayward, K. Parmesar, M. A. Saleem, What is circulating factor disease and how is it currently explained? Pediatr Nephrol 38, 3513–3518 (2023).

18. R. Menon, E. A. Otto, P. Hoover, S. Eddy, L. Mariani, B. Godfrey, C. C. Berthier, F. Eichinger, L. Subramanian, J. Harder, W. Ju, V. Nair, M. Larkina, A. S. Naik, J. Luo, S. Jain, R. Sealfon, O. Troyanskaya, N. Hacohen, J. B. Hodgin, M. Kretzler, K. P. M. Project (KPMP), N. S. S. Network (NEPTUNE), Single cell transcriptomics identifies focal segmental glomerulosclerosis remission endothelial biomarker (2020), doi:10.1172/jci.insight.133267.

19. L. H. Mariani, S. Eddy, F. M. AlAkwaa, P. J. McCown, J. L. Harder, V. Nair, F. Eichinger, S. Martini, A. D. Ademola, V. Boima, H. N. Reich, J. El Saghir, B. Godfrey, W. Ju, E. C. Tanner, V. Vega-Warner, N. L. Wys, S. G. Adler, G. B. Appel, A. Athavale, M. A. Atkinson, S. M. Bagnasco, L. Barisoni, E. Brown, D. C. Cattran, G. M. Coppock, K. M. Dell, V. K. Derebail, F. C. Fervenza, A. Fornoni, C. A. Gadegbeku, K. L. Gibson, L. A. Greenbaum, S. R. Hingorani, M. A. Hladunewich, J. B. Hodgin, M. C. Hogan, L. B. Holzman, J. A. Jefferson, F. J. Kaskel, J. B. Kopp, R. A. Lafayette, K. V. Lemley, J. C. Lieske, J.-J. Lin, R. Menon, K. E. Meyers, P. H. Nachman, C. C. Nast, M. M. O’Shaughnessy, E. A. Otto, K. J. Reidy, K. K. Sambandam, J. R. Sedor, C. B. Sethna, P. Singer, T. Srivastava, C. L. Tran, K. R. Tuttle, S. M. Vento, C.-S. Wang, A. O. Ojo, D. Adu, D. S. Gipson, H. Trachtman, M. Kretzler, Precision nephrology identified tumor necrosis factor activation variability in minimal change disease and focal segmental glomerulosclerosis. Kidney Int 103, 565–579 (2023).

20. K. Z. Latt, J. Heymann, J. H. Jessee, A. Z. Rosenberg, C. C. Berthier, A. Arazi, S. Eddy, T. Yoshida, Y. Zhao, V. Chen, G. W. Nelson, M. Cam, P. Kumar, M. Mehta, M. C. Kelly, M. Kretzler, P. E. Ray, M. Moxey-Mims, G. H. Gorman, B. L. Lechner, R. Regunathan-Shenk, D. S. Raj, K. Susztak, C. A. Winkler, J. B. Kopp, Urine Single-Cell RNA Sequencing in Focal Segmental Glomerulosclerosis Reveals Inflammatory Signatures. Kidney International Reports 7, 289–304 (2022).

21. J. M. McDaniels, A. C. Shetty, T. V. Rousselle, E. Bardhi, D. G. Maluf, V. R. Mas, The cellular landscape of the normal kidney allograft: Main players balancing the alloimmune response. Frontiers in Transplantation 1 (2022) (available at https://www.frontiersin.org/articles/10.3389/frtra.2022.988238).

22. J. M. McDaniels, A. C. Shetty, C. Kuscu, C. Kuscu, E. Bardhi, T. Rousselle, C. Drachenberg, M. Talwar, J. D. Eason, T. Muthukumar, D. G. Maluf, V. R. Mas, Single nuclei transcriptomics delineates complex immune and kidney cell interactions contributing to kidney allograft fibrosis. Kidney Int 103, 1077–1092 (2023).

23. Y. Muto, P. C. Wilson, N. Ledru, H. Wu, H. Dimke, S. S. Waikar, B. D. Humphreys, Single cell transcriptional and chromatin accessibility profiling redefine cellular heterogeneity in the adult human kidney. Nat Commun 12, 2190 (2021).

24. M. T. Lindenmeyer, F. Eichinger, K. Sen, H.-J. Anders, I. Edenhofer, D. Mattinzoli, M. Kretzler, M. P. Rastaldi, C. D. Cohen, Systematic Analysis of a Novel Human Renal Glomerulus-Enriched Gene Expression Dataset. PLOS ONE 5, e11545 (2010).

25. E. B. Peixoto, A. Papadimitriou, D. a. T. Teixeira, C. Montemurro, D. A. Duarte, K. C. Silva, P. P. Joazeiro, J. M. Lopes de Faria, J. B. Lopes de Faria, Reduced LRP6 expression and increase in the interaction of GSK3β with p53 contribute to podocyte apoptosis in diabetes mellitus and are prevented by green tea. J Nutr Biochem 26, 416– 430 (2015).

26. G. Caridi, F. Perfumo, G. M. Ghiggeri, NPHS2 (Podocin) Mutations in Nephrotic Syndrome. Clinical Spectrum and Fine Mechanisms. Pediatr Res 57, 54–61 (2005).

27. M. Hu, R. Wang, X. Li, M. Fan, J. Lin, J. Zhen, L. Chen, Z. Lv, LncRNA MALAT1 is dysregulated in diabetic nephropathy and involved in high glucose-induced podocyte injury via its interplay with β-catenin. J Cell Mol Med 21, 2732–2747 (2017).

28. X. Li, I. Venkatesh, V. Villanueva, H. Wei, T. Geraghty, A. Rajagopalan, R. W. Helmuth, M. M. Altintas, H. M. Faridi, V. Gupta, Podocyte-specific deletion of miR-146a increases podocyte injury and diabetic kidney disease. Front. Med. 9 (2022), doi:10.3389/fmed.2022.897188.

29. Y. Sun, M. Deng, X. Ke, X. Lei, H. Ju, Z. Liu, X. Bai, Epidermal Growth Factor Protects Against High Glucose-Induced Podocyte Injury Possibly via Modulation of Autophagy and PI3K/AKT/mTOR Signaling Pathway Through DNA Methylation. DMSO 14, 2255–2268 (2021).

30. J. Hong, X. Li, Y. Hao, H. Xu, L. Yu, Z. Meng, J. Zhang, M. Zhu, The PRMT6/STAT1/ACSL1 axis promotes ferroptosis in diabetic nephropathy. Cell Death Differ, 1–15 (2024).

31. N. Song, F. Thaiss, L. Guo, NFκB and Kidney Injury. Front. Immunol. 10 (2019), doi:10.3389/fimmu.2019.00815.

32. H. Wang, Y. Zhang, F. Xia, W. Zhang, P. Chen, G. Yang, Protective effect of silencing Stat1 on high glucose-induced podocytes injury via Forkhead transcription factor O1-regulated the oxidative stress response. BMC Molecular and Cell Biology 20, 27 (2019).

33. A. Staruschenko, O. Palygin, D. V. Ilatovskaya, T. S. Pavlov, Epidermal growth factors in the kidney and relationship to hypertension. American Journal of Physiology-Renal Physiology 305, F12–F20 (2013).

34. G. Bollée, M. Flamant, S. Schordan, C. Fligny, E. Rumpel, M. Milon, E. Schordan, N. Sabaa, S. Vandermeersch, A. Galaup, A. Rodenas, I. Casal, S. W. Sunnarborg, D. J. Salant, J. B. Kopp, D. W. Threadgill, S. E. Quaggin, J.-C. Dussaule, S. Germain, L. Mesnard, K. Endlich, C. Boucheix, X. Belenfant, P. Callard, N. Endlich, P.-L. Tharaux, The Epidermal Growth Factor Receptor Promotes Glomerular Injury and Renal Failure in Rapidly Progressive Crescentic Glomerulonephritis; the Identification of Possible Therapy. Nat Med 17, 1242–1250 (2011).

35. L. R. Harskamp, R. T. Gansevoort, H. van Goor, E. Meijer, The epidermal growth factor receptor pathway in chronic kidney diseases. Nat Rev Nephrol 12, 496–506 (2016).

36. K. Datta, J. Li, S. A. Karumanchi, E. Wang, E. Rondeau, D. Mukhopadhyay, Regulation of vascular permeability factor/vascular endothelial growth factor (VPF/VEGF-A) expression in podocytes. Kidney International 66, 1471–1478 (2004).

37. G. A. Sivaskandarajah, M. Jeansson, Y. Maezawa, V. Eremina, H. J. Baelde, S. E. Quaggin, Vegfa protects the glomerular microvasculature in diabetes. Diabetes 61, 2958–2966 (2012).

38. D. Veron, P. K. Aggarwal, Q. Li, G. Moeckel, M. Kashgarian, A. Tufro, Podocyte VEGF-A Knockdown Induces Diffuse Glomerulosclerosis in Diabetic and in eNOS Knockout Mice. Front Pharmacol 12, 788886 (2022).

39. D. Veron, K. J. Reidy, C. Bertuccio, J. Teichman, G. Villegas, J. Jimenez, W. Shen, J. B. Kopp, D. B. Thomas, A. Tufro, Overexpression of VEGF-A in podocytes of adult mice causes glomerular disease. Kidney Int 77, 989–999 (2010).

40. A. Tufro, D. Veron, VEGF AND PODOCYTES IN DIABETIC NEPHROPATHY. Semin Nephrol 32, 385–393 (2012).

41. T. Okamoto, S. Sasaki, T. Yamazaki, Y. Sato, H. Ito, T. Ariga, Prevalence of CD44-positive glomerular parietal epithelial cells reflects podocyte injury in adriamycin nephropathy. Nephron Exp Nephrol 124, 11–18 (2013).

42. M. Nagata, Podocyte injury and its consequences. Kidney International 89, 1221–1230 (2016).

43. N. Ito, K. Sakamoto, C. Hikichi, T. Matsusaka, M. Nagata, Biphasic MIF and SDF1 expression during podocyte injury promote CD44-mediated glomerular parietal cell migration in focal segmental glomerulosclerosis. American Journal of Physiology-Renal Physiology 318, F741–F753 (2020).

44. N. Chebotareva, A. Vinogradov, L. Tsoy, V. Varshavskiy, E. Stoljarevich, A. Bugrova, Y. Lerner, T. Krasnova, E. Biryukova, A. S. Kononikhin, CD44 Expression in Renal Tissue Is Associated with an Increase in Urinary Levels of Complement Components in Chronic Glomerulopathies. Int J Mol Sci 24, 7190 (2023).

45. V. D. D’Agati, S. J. Shankland, Recognizing diversity in parietal epithelial cells. Kidney International 96, 16–19 (2019).

46. C. Sagrinati, G. S. Netti, B. Mazzinghi, E. Lazzeri, F. Liotta, F. Frosali, E. Ronconi, C. Meini, M. Gacci, R. Squecco, M. Carini, L. Gesualdo, F. Francini, E. Maggi, F. Annunziato, L. Lasagni, M. Serio, S. Romagnani, P. Romagnani, Isolation and characterization of multipotent progenitor cells from the Bowman’s capsule of adult human kidneys. J Am Soc Nephrol 17, 2443–2456 (2006).

47. P. Romagnani, Toward the Identification of a “Renopoietic System”? Stem Cells 27, 2247–2253 (2009).

48. K. K. Ratnam, X. Feng, P. Y. Chuang, V. Verma, T.-C. Lu, J. Wang, Y. Jin, E. F. Farias, J. L. Napoli, N. Chen, L. Kaufman, T. Takano, V. D. D’Agati, P. E. Klotman, J. C. He, Role of the retinoic acid receptor-α in HIV-associated nephropathy. Kidney Int 79, 624–634 (2011).

49. B. He, P. Chen, S. Zambrano, D. Dabaghie, Y. Hu, K. Möller-Hackbarth, D. Unnersjö-Jess, G. G. Korkut, E. Charrin, M. Jeansson, M. Bintanel-Morcillo, A. Witasp, L. Wennberg, A. Wernerson, B. Schermer, T. Benzing, P. Ernfors, C. Betsholtz, M. Lal, R. Sandberg, J. Patrakka, Single-cell RNA sequencing reveals the mesangial identity and species diversity of glomerular cell transcriptomes. Nat Commun 12, 2141 (2021).

50. B. B. Lake, R. Menon, S. Winfree, Q. Hu, R. Melo Ferreira, K. Kalhor, D. Barwinska, E. A. Otto, M. Ferkowicz, D. Diep, N. Plongthongkum, A. Knoten, S. Urata, L. H. Mariani, A. S. Naik, S. Eddy, B. Zhang, Y. Wu, D. Salamon, J. C. Williams, X. Wang, K. S. Balderrama, P. J. Hoover, E. Murray, J. L. Marshall, T. Noel, A. Vijayan, A. Hartman, F. Chen, S. S. Waikar, S. E. Rosas, F. P. Wilson, P. M. Palevsky, K. Kiryluk, J. R. Sedor, R. D. Toto, C. R. Parikh, E. H. Kim, R. Satija, A. Greka, E. Z. Macosko, P. V. Kharchenko, J. P. Gaut, J. B. Hodgin, KPMP Consortium, M. T. Eadon, P. C. Dagher, T. M. El-Achkar, K. Zhang, M. Kretzler, S. Jain, An atlas of healthy and injured cell states and niches in the human kidney. Nature 619, 585–594 (2023).

51. A. Bottos, E. Destro, A. Rissone, S. Graziano, G. Cordara, B. Assenzio, M. R. Cera, L. Mascia, F. Bussolino, M. Arese, The synaptic proteins neurexins and neuroligins are widely expressed in the vascular system and contribute to its functions. Proc Natl Acad Sci U S A 106, 20782–20787 (2009).

52. X. Gu, S. Zhang, T. Zhang, Abnormal Crosstalk between Endothelial Cells and Podocytes Mediates Tyrosine Kinase Inhibitor (TKI)-Induced Nephrotoxicity. Cells 10, 869 (2021).

53. J. J. Olsen, S. Ö.-G. Pohl, A. Deshmukh, M. Visweswaran, N. C. Ward, F. Arfuso, M. Agostino, A. Dharmarajan, The Role of Wnt Signalling in Angiogenesis. Clin Biochem Rev 38, 131–142 (2017).

54. H. Oberleithner, C. Riethmüller, H. Schillers, G. A. MacGregor, H. E. de Wardener, M. Hausberg, Plasma sodium stiffens vascular endothelium and reduces nitric oxide release. Proc Natl Acad Sci U S A 104, 16281–16286 (2007).

55. H. Oberleithner, M. Wälte, K. Kusche-Vihrog, Sodium renders endothelial cells sticky for red blood cells. Front Physiol 6, 188 (2015).

56. P. E. Westerweel, M. C. Verhaar, Protective Actions of PPAR-γ Activation in Renal Endothelium. PPAR Res 2008, 635680 (2008).

57. J. Wollborn, S. Siemering, C. Steiger, H. Buerkle, U. Goebel, M. A. Schick, Phosphodiesterase-4 inhibition reduces ECLS-induced vascular permeability and improves microcirculation in a rodent model of extracorporeal resuscitation. American Journal of Physiology-Heart and Circulatory Physiology 316, H751–H761 (2019).

58. J. Surapisitchat, K.-I. Jeon, C. Yan, J. A. Beavo, Differential Regulation of Endothelial Cell Permeability by cGMP via Phosphodiesterases 2 and 3. Circulation Research 101, 811–818 (2007).

59. Y.-C. Lin, H. Samardzic, R. H. Adamson, E. M. Renkin, J. F. Clark, R. K. Reed, F.-R. E. Curry, Phosphodiesterase 4 inhibition attenuates atrial natriuretic peptide-induced vascular hyperpermeability and loss of plasma volume. The Journal of Physiology 589, 341–353 (2011).

60. Y. Kirita, H. Wu, K. Uchimura, P. C. Wilson, B. D. Humphreys, Cell profiling of mouse acute kidney injury reveals conserved cellular responses to injury. Proceedings of the National Academy of Sciences 117, 15874–15883 (2020).

61. S. Bruno, B. Bussolati, C. Grange, F. Collino, M. E. Graziano, U. Ferrando, G. Camussi, CD133+ renal progenitor cells contribute to tumor angiogenesis. Am J Pathol 169, 2223–2235 (2006).

62. E. Ronconi, C. Sagrinati, M. L. Angelotti, E. Lazzeri, B. Mazzinghi, L. Ballerini, E. Parente, F. Becherucci, M. Gacci, M. Carini, E. Maggi, M. Serio, G. B. Vannelli, L. Lasagni, S. Romagnani, P. Romagnani, Regeneration of glomerular podocytes by human renal progenitors. J Am Soc Nephrol 20, 322–332 (2009).

63. Y. Chen, Y. Gong, J. Zou, G. Li, F. Zhang, Y. Yang, Y. Liang, W. Dai, L. He, H. Lu, Single-cell transcriptomic analysis reveals transcript enrichment in oxidative phosphorylation, fluid sheer stress, and inflammatory pathways in obesity-related glomerulopathy. Genes Dis 11, 101101 (2024).

64. D. Deleersnijder, A. H. Van Craenenbroeck, B. Sprangers, Deconvolution of Focal Segmental Glomerulosclerosis Pathophysiology Using Transcriptomics Techniques. Glomerular Dis 1, 265–276 (2021).

65. D. Roccatello, A. Baffa, C. Naretto, A. Barreca, R. Cravero, E. Roscini, S. Sciascia, R. Fenoglio, Focal segmental glomerular sclerosis can be effectively treated using an intensive B-cell depletion therapy. Clinical Kidney Journal 16, 1258–1264 (2023).

66. A. D. Morris, L. Floyd, A. Woywodt, A. Dhaygude, Rituximab in the treatment of primary FSGS: time for its use in routine clinical practice? Clin Kidney J 16, 1199–1205 (2023).

67. M. Colucci, J. Oniszczuk, M. Vivarelli, V. Audard, B-Cell Dysregulation in Idiopathic Nephrotic Syndrome: What We Know and What We Need to Discover. Front. Immunol. 13 (2022), doi:10.3389/fimmu.2022.823204.

68. S. D. Ricardo, H. van Goor, A. A. Eddy, Macrophage diversity in renal injury and repair. J Clin Invest 118, 3522–3530 (2008).

69. R. K. Tadagavadi, W. B. Reeves, Renal Dendritic Cells Ameliorate Nephrotoxic Acute Kidney Injury. J Am Soc Nephrol 21, 53–63 (2010).

70. Q. Cao, J. Lu, Q. Li, C. Wang, X. M. Wang, V. W. S. Lee, C. Wang, H. Nguyen, G. Zheng, Y. Zhao, S. I. Alexander, Y. Wang, D. C. H. Harris, CD103+ Dendritic Cells Elicit CD8+ T Cell Responses to Accelerate Kidney Injury in Adriamycin Nephropathy. J Am Soc Nephrol 27, 1344–1360 (2016).

71. R. Wang, T. Chen, C. Wang, Z. Zhang, X. M. Wang, Q. Li, V. W. S. Lee, Y. M. Wang, G. Zheng, S. I. Alexander, Y. Wang, D. C. H. Harris, Q. Cao, Flt3 inhibition alleviates chronic kidney disease by suppressing CD103+ dendritic cell-mediated T cell activation. Nephrol Dial Transplant 34, 1853–1863 (2019).

72. S. Li, S. Jiang, Q. Zhang, B. Jin, D. Lv, W. Li, M. Zhao, C. Jiang, C. Dai, Z. Liu, Integrin β3 Induction Promotes Tubular Cell Senescence and Kidney Fibrosis. Front Cell Dev Biol 9, 733831 (2021).

73. V. Rapisarda, M. Borghesan, V. Miguela, V. Encheva, A. P. Snijders, A. Lujambio, A. O’Loghlen, Integrin Beta 3 Regulates Cellular Senescence by Activating the TGF-β Pathway. Cell Rep 18, 2480–2493 (2017).

74. J. Zhao, X. Jiang, L. Yan, J. Lin, H. Guo, S. Yu, B. Ye, J. Zhu, W. Zhang, Retinoic acid inducible gene-I slows down cellular senescence through negatively regulating the integrin β3/p38 MAPK pathway. Cell Cycle 18, 3378– 3392 (2019).

75. D. Zhou, R. J. Tan, L. Lin, L. Zhou, Y. Liu, Activation of hepatocyte growth factor receptor, c-met, in renal tubules is required for renoprotection after acute kidney injury. Kidney Int 84, 509–520 (2013).

76. P. A. Agustian, M. Schiffer, W. Gwinner, I. Schäfer, K. Theophile, F. Modde, C. L. Bockmeyer, J. Traeder, U. Lehmann, A. Grosshennig, H. H. Kreipe, V. Bröcker, J. U. Becker, Diminished met signaling in podocytes contributes to the development of podocytopenia in transplant glomerulopathy. Am J Pathol 178, 2007–2019 (2011).

77. J. Shoji, A. Mii, M. Terasaki, A. Shimizu, Update on Recurrent Focal Segmental Glomerulosclerosis in Kidney Transplantation. NEF 144, 65–70 (2020).

78. G. Salfi, F. Casiraghi, G. Remuzzi, Current understanding of the molecular mechanisms of circulating permeability factor in focal segmental glomerulosclerosis. Front Immunol 14, 1247606 (2023).

79. L. Gallon, J. Leventhal, A. Skaro, Y. Kanwar, A. Alvarado, Resolution of recurrent focal segmental glomerulosclerosis after retransplantation. N Engl J Med 366, 1648–1649 (2012).

80. H. R. Al Shamsi, I. Shaheen, D. Aziz, Management of recurrent focal segmental glomerulosclerosis (FSGS) post renal transplantation. Transplant Rev (Orlando) 36, 100675 (2022).

81. A. S. De Vriese, J. F. Wetzels, R. J. Glassock, S. Sethi, F. C. Fervenza, Therapeutic trials in adult FSGS: lessons learned and the road forward. Nat Rev Nephrol 17, 619–630 (2021).

82. P. T. Brinkkoetter, T. Bork, S. Salou, W. Liang, A. Mizi, C. Özel, S. Koehler, H. H. Hagmann, C. Ising, A. Kuczkowski, S. Schnyder, A. Abed, B. Schermer, T. Benzing, O. Kretz, V. G. Puelles, S. Lagies, M. Schlimpert, B. Kammerer, C. Handschin, C. Schell, T. B. Huber, Anaerobic Glycolysis Maintains the Glomerular Filtration Barrier Independent of Mitochondrial Metabolism and Dynamics. Cell Rep 27, 1551–1566.e5 (2019).

83. S. Ozawa, S. Ueda, H. Imamura, K. Mori, K. Asanuma, M. Yanagita, T. Nakagawa, Glycolysis, but not Mitochondria, responsible for intracellular ATP distribution in cortical area of podocytes. Sci Rep 5, 18575 (2015).

84. A. G. Carrasco, A. Izquierdo-Lahuerta, Á. M. Valverde, L. Ni, E. Flores-Salguero, R. J. Coward, G. Medina-Gómez, The protective role of peroxisome proliferator-activated receptor gamma in lipotoxic podocytes. Biochim Biophys Acta Mol Cell Biol Lipids 1868, 159329 (2023).

85. A. Fornoni, S. Merscher, Lipid Metabolism Gets in a JAML during Kidney Disease. Cell Metab 32, 903–905 (2020).

86. T. Kabashima, T. Kawaguchi, B. E. Wadzinski, K. Uyeda, Xylulose 5-phosphate mediates glucose-induced lipogenesis by xylulose 5-phosphate-activated protein phosphatase in rat liver. Proc Natl Acad Sci U S A 100, 5107– 5112 (2003).

87. Y. Q. Liu, K. Uyeda, A mechanism for fatty acid inhibition of glucose utilization in liver. Role of xylulose 5-P. J Biol Chem 271, 8824–8830 (1996).

88. J. A. Schaub, F. M. AlAkwaa, P. J. McCown, A. S. Naik, V. Nair, S. Eddy, R. Menon, E. A. Otto, J. Hartman, D. Fermin, C. O’Connor, M. Bitzer, R. Harned, P. Ladd, L. Pyle, J. B. Hodgin, F. C. Brosius, R. G. Nelson, M. Kretzler, P. Bjornstad, SGLT2 inhibition mitigates perturbations in nephron segment-specific metabolic transcripts and mTOR pathway activity in kidneys of young persons with type 2 diabetes, 2022.07.23.22277943 (2022).

89. Z.-W. Dai, K.-D. Cai, L.-C. Xu, L.-L. Wang, Perilipin2 inhibits diabetic nephropathy-induced podocyte apoptosis by activating the PPARγ signaling pathway. Molecular and Cellular Probes 53, 101584 (2020).

90. Q. Chen, H. Jiang, R. Ding, J. Zhong, L. Li, J. Wan, X. Feng, L. Peng, X. Yang, H. Chen, A. Wang, J. Jiao, Q. Yang, X. Chen, X. Li, L. Shi, G. Zhang, M. Wang, H. Yang, Q. Li, Cell-type-specific molecular characterization of cells from circulation and kidney in IgA nephropathy with nephrotic syndrome. Front Immunol 14, 1231937 (2023).

91. M. R. Pollak, Familial FSGS. Adv Chronic Kidney Dis 21, 422–425 (2014).

92. J.-J. Chung, L. Goldstein, Y.-J. J. Chen, J. Lee, J. D. Webster, M. Roose-Girma, S. C. Paudyal, Z. Modrusan, A. Dey, A. S. Shaw, Single-Cell Transcriptome Profiling of the Kidney Glomerulus Identifies Key Cell Types and Reactions to Injury. J Am Soc Nephrol 31, 2341–2354 (2020).

93. S. Liu, Y. Zhao, S. Lu, T. Zhang, M. T. Lindenmeyer, V. Nair, S. E. Gies, G. Wu, R. G. Nelson, J. Czogalla, H. Aypek, S. Zielinski, Z. Liao, M. Schaper, D. Fermin, C. D. Cohen, D. Delic, C. F. Krebs, F. Grahammer, T. Wiech, M. Kretzler, C. Meyer-Schwesinger, S. Bonn, T. B. Huber, Single-cell transcriptomics reveals a mechanosensitive injury signaling pathway in early diabetic nephropathy. Genome Med 15, 2 (2023).

94. S.-Y. Li, J. Park, C. Qiu, S. H. Han, M. B. Palmer, Z. Arany, K. Susztak, Increasing the level of peroxisome proliferator-activated receptor **γ** coactivator-1**α** in podocytes results in collapsing glomerulopathy. JCI Insight 2 (2017), doi:10.1172/jci.insight.92930.

95. Y.-C. Tu, H.-P. Shu, L.-L. Sun, Q.-Q. Liao, L. Feng, M. Ren, L.-J. Yao, The Physiopathologic Roles of Calcium Signaling in Podocytes. FBL 28, 240 (2023).

96. S. Yu, W.-I. Choi, Y. J. Choi, H.-Y. Kim, F. Hildebrandt, H. Y. Gee, PLCE1 regulates the migration, proliferation, and differentiation of podocytes. Exp Mol Med 52, 594–603 (2020).

97. M. Trebak, L. Lemonnier, J. T. Smyth, G. Vazquez, J. W. Putney, in Transient Receptor Potential (TRP) Channels, V. Flockerzi, B. Nilius, Eds. (Springer, Berlin, Heidelberg, 2007), pp. 593–614.

98. X. Yang, D. Wu, H. Du, F. Nie, X. Pang, Y. Xu, MicroRNA-135a is involved in podocyte injury in a transient receptor potential channel 1-dependent manner. Int J Mol Med 40, 1511–1519 (2017).

99. T. Struk, V. Nair, F. Eichinger, M. Kretzler, R. Wedlich-Söldner, S. Bayraktar, H. Pavenstädt, Transcriptome analysis of primary podocytes reveals novel calcium regulated regulatory networks. The FASEB Journal 34, 14490– 14506 (2020).

100. L. Zhang, T. Ji, Q. Wang, K. Meng, R. Zhang, H. Yang, C. Liao, L. Ma, J. Jiao, Calcium-Sensing Receptor Stimulation in Cultured Glomerular Podocytes Induces TRPC6-Dependent Calcium Entry and RhoA Activation. Cell Physiol Biochem 43, 1777–1789 (2017).

101. N. Hempel, M. Trebak, Crosstalk between Calcium and Reactive Oxygen Species Signaling in Cancer. Cell Calcium 63, 70–96 (2017).

102. C. G. Economou, P. V. Kitsiou, A. K. Tzinia, E. Panagopoulou, E. Marinos, D. B. Kershaw, D. Kerjaschki, E. C. Tsilibary, Enhanced podocalyxin expression alters the structure of podocyte basal surface. J Cell Sci 117, 3281– 3294 (2004).

103. M. Hara, K. Yamagata, Y. Tomino, A. Saito, Y. Hirayama, S. Ogasawara, H. Kurosawa, S. Sekine, K. Yan, Urinary podocalyxin is an early marker for podocyte injury in patients with diabetes: establishment of a highly sensitive ELISA to detect urinary podocalyxin. Diabetologia 55, 2913–2919 (2012).

104. M. W. Seiler, H. G. Rennke, M. A. Venkatachalam, R. S. Cotran, Pathogenesis of polycation-induced alterations (“fusion”) of glomerular epithelium. Lab Invest 36, 48–61 (1977).

105. T. Takeda, W. Y. Go, R. A. Orlando, M. G. Farquhar, Expression of podocalyxin inhibits cell-cell adhesion and modifies junctional properties in Madin-Darby canine kidney cells. Mol Biol Cell 11, 3219–3232 (2000).

106. D. Appel, D. B. Kershaw, B. Smeets, G. Yuan, A. Fuss, B. Frye, M. Elger, W. Kriz, J. Floege, M. J. Moeller, Recruitment of podocytes from glomerular parietal epithelial cells. J Am Soc Nephrol 20, 333–343 (2009).

107. D. G. Eng, M. W. Sunseri, N. V. Kaverina, S. S. Roeder, J. W. Pippin, S. J. Shankland, Glomerular parietal epithelial cells contribute to adult podocyte regeneration in experimental focal segmental glomerulosclerosis. Kidney Int 88, 999–1012 (2015).

108. L. Miesen, E. Steenbergen, B. Smeets, Parietal cells—new perspectives in glomerular disease. Cell Tissue Res 369, 237–244 (2017).

109. J. C. Chandler, D. J. Jafree, S. Malik, G. Pomeranz, M. Ball, M. Kolatsi-Joannou, A. Piapi, W. J. Mason, A. V. Benest, D. O. Bates, A. Letunovska, R. Al-Saadi, M. Rabant, O. Boyer, K. Pritchard-Jones, P. J. Winyard, A. S. Mason, A. S. Woolf, A. M. Waters, D. A. Long, Single-cell transcriptomics identifies aberrant glomerular angiogenic signalling in the early stages of WT1 kidney disease. J Pathol 264, 212–227 (2024).

110. L. Gallon, H. Zubair, A. C. Shetty, J. McDaniels, T. V. Rousselle, S. Azim, C. Kuscu, C. Kuscu, J. D. Eason, D. Maluf, V. Mas, Changes in Endothelial and Proximal Tubule Cells in Focal Segmental Glomerulosclerosis: Single-Cell Resolution of Human Renal Allografts with Recurrent Disease (available at https://atc.digitellinc.com/p/s/changes-in-endothelial-and-proximal-tubule-cells-in-focal-segmental-glomerulosclerosis-single-cell-resolution-of-human-renal-allografts-with-recurrent-disease-35604).

111. S. Zambrano, L. He, T. Kano, Y. Sun, E. Charrin, M. Lal, C. Betsholtz, Y. Suzuki, J. Patrakka, Molecular insights into the early stage of glomerular injury in IgA nephropathy using single-cell RNA sequencing. Kidney International 101, 752–765 (2022).

112. J. Patrakka, V. Ruotsalainen, P. Reponen, E. Qvist, J. Laine, C. Holmberg, K. Tryggvason, H. Jalanko, RECURRENCE OF NEPHROTIC SYNDROME IN KIDNEY GRAFTS OF PATIENTS WITH CONGENITAL NEPHROTIC SYNDROME OF THE FINNISH TYPE: Role of Nephrin: 1. Transplantation 73, 394 (2002).

113. D. S. Gipson, C.-S. Wang, E. Salmon, R. Gbadegesin, A. Naik, S. Sanna-Cherchi, A. Fornoni, M. Kretzler, S. Merscher, P. Hoover, K. Kidwell, M. Saleem, L. Riella, L. Holzman, A. Jackson, O. Olabisi, P. Cravedi, B. S. Freedman, J. Himmelfarb, M. Vivarelli, J. Harder, J. Klein, G. Burke, M. Rheault, C. Spino, H. E. Desmond, H. Trachtman, FSGS Recurrence Collaboration: Report of a Symposium. Glomerular Dis 4, 1–10 (2023).

114. P. C. Wilson, H. Wu, Y. Kirita, K. Uchimura, N. Ledru, H. G. Rennke, P. A. Welling, S. S. Waikar, B. D. Humphreys, The single-cell transcriptomic landscape of early human diabetic nephropathy. Proc Natl Acad Sci U S A 116, 19619–19625 (2019).

