## Supplementary Tables (S1-S4) and Figures (S1-S9) for "Cellular and Molecular Resolution of Focal Segmental Glomerulosclerosis Recurrence in Human Allografts"

**Table S1: Summary of Illumina sequencing results.**

| Sample Type | | Sample | Total number of nuclei | Mean reads per cells | Median genes per cell | Total number of reads | Total genes detected | % of reads mapped to the genome | % reads mapped to exons | % reads mapped to intergenic |
|---|---|---|---|---|---|---|---|---|---|---|
| **Native** | Normal-1 | GSM3823939 (nNK1) | | | | | | | | |
| | Normal-2 | GSM3823940 (nNK2) | | | | | | | | |
| | Normal-3 | GSM3823941 (nNK3) | | | | | | | | |
| | FSGS | NW-3 (nFSGS) | 5,624 | 72,499 | 1,297 | 407,736,246 | 26,258 | 38.6 | 35.1 | 3.8 |
| **Allograft** | Normal-1 | NW-4 (NKTx1) | 8,520 | 69,376 | 1,556 | 591,080,967 | 28,736 | 45.5 | 41.6 | 4.3 |
| | Normal-2 | KUT023 (NKTx2) | 8,077 | 70,157 | 1,661 | 566,655,067 | 30,556 | 76.5 | 53.9 | 5.6 |
| | Normal-3 | KUT060 (NKTx3) | 7,495 | 135,222 | 1,596 | 1,013,490,932 | 29,254 | 77.3 | 40.4 | 5.3 |
| | Normal-4 | KUT081 (NKTx4) | 4,634 | 134,795 | 845 | 624,642,131 | 29,126 | 70.8 | 33.3 | 4.1 |
| | FSGS-T1 | NW-1 (KTxFSGS1-T1) | 2,669 | 157,882 | 904 | 421,386,244 | 24,150 | 63.7 | 21.1 | 3.2 |

| | | | | | | | | | | |
|---|---|---|---|---|---|---|---|---|---|---|
| | FSGS-T2 | NW-2 (KTxFSGS1-T2) | 5,769 | 69,657 | 1,809 | 401,853,270 | 26,678 | 70.3 | 43.2 | 4.6 |
| | FSGS-T1 | NW-5 (KTxFSGS2-T1) | 3,116 | 126,065 | 1,352 | 399,121,180 | 25,644 | 32.3 | 29.3 | 3.3 |
| | FSGS-T1 | NW-6 (KTxFSGS3-T1) | 6,858 | 78,708 | 1,099 | 539,776,539 | 27,728 | 25.9 | 24.3 | 2.7 |
| | FSGS-T2 | NW-7 (KTxFSGS2-T2) | 313 | 1,236,705 | 2,201 | 387,088,730 | 22,030 | 72.9 | 68.8 | 4.8 |

The summary metrics for all samples are presented before further downstream analysis. nNK: native normal kidney, NKTx: normal allograft, nFSGS: native kidney with FSGS, KTxFSGS-T1: kidney allograft with first recurrence of FSGS post-kidney transplantation, KTxFSGS-T2: kidney allograft with second recurrence of FSGS post- kidney transplantation.

**Table S2. Summary of Q30 scores.**

| Sample Type | | Sample | Q30 Bases in percentage | | | |
|---|---|---|---|---|---|---|
| | | | Barcode | RNA Read 1 | RNA Read 2 | UMI |
| Native | nNK1 | GSM3823939 | | | | |
| | nNK2 | GSM3823940 | | | | |
| | nNK3 | GSM3823941 | | | | |
| | nFSGS | NW-3 | 97.0 | 93.9 | 81.6 | 96.7 |
| Allograft | NKTx1 | NW-4 | 97.0 | 93.8 | 78.6 | 96.8 |
| | NKTx2 | KUT023 | 97.2 | 93.7 | 84.9 | 96.8 |
| | NKTx3 | KUT060 | 97.1 | 93.7 | 85.3 | 94.8 |
| | NKTx4 | KUT081 | 97.1 | 93.7 | 82.6 | 96.8 |
| | KTxFSGS1-T1 | NW-1 | 96.9 | 94.2 | 79.0 | 96.6 |
| | KTxFSGS1-T2 | NW-2 | 97.0 | 94.3 | 81.9 | 96.7 |
| | KTxFSGS2-T1 | NW-5 | 97.0 | 93.9 | 80.2 | 96.7 |
| | KTxFSGS3-T1 | NW-6 | 96.9 | 93.0 | 73.6 | 96.7 |
| | KTxFSGS2-T2 | NW-7 | 97.1 | 93.3 | 85.5 | 96.8 |

UMI, unique molecular identifier, nNK: native normal kidney, NKTx: normal allograft, nFSGS: native kidney with FSGS, KTxFSGS-T1: kidney allograft with first recurrence of FSGS post-kidney transplantation, KTxFSGS-T2: kidney allograft with second recurrence of FSGS post- kidney transplantation.

**Table S3. Counts of isolated cells in each cell cluster.**

| **Cell Cluster** | **nNK** | **NKTx** | **nFSGS** | **KTxFSGS1** | **KTxFSGS2** | **Total** |
|---|---|---|---|---|---|---|
| Proximal Tubule 1 (PT-1) | 3339 | 5877 | 1255 | 2091 | 905 | 13467 |
| Proximal Tubule 2 (PT-2) | 452 | 935 | 832 | 1320 | 900 | 4439 |
| Distal Tubule 1 (DT-1) | 1978 | 5089 | 1095 | 1792 | 1371 | 11325 |
| Distal Tubule 2 (DT-2) | 1847 | 2160 | 164 | 745 | 145 | 5061 |
| Endothelial 1 (EC-1) | 369 | 1755 | 232 | 799 | 256 | 3411 |
| Endothelial 2 (EC-2) | 147 | 806 | 64 | 343 | 70 | 1430 |
| Endothelial 3 (EC-3) | 6 | 74 | 15 | 43 | 16 | 154 |
| Collecting Duct Principal 1 (CDP1) | 1072 | 1836 | 307 | 1122 | 223 | 4560 |
| Collecting Duct Principal 2 (CDP2) | 932 | 1610 | 309 | 842 | 510 | 4203 |
| Collecting Duct Intercalated 1 (CDI1) | 1158 | 1628 | 147 | 508 | 99 | 3540 |
| Collecting Duct Intercalated 2 (CDI2) | 266 | 719 | 18 | 175 | 33 | 1211 |
| Podocyte (POD) | 366 | 598 | 86 | 450 | 41 | 1541 |
| Parietal Epithelial (PEC) | 391 | 479 | 137 | 347 | 185 | 1539 |
| Immune (IMM) | 33 | 569 | 146 | 495 | 291 | 1534 |
| Unknown 1 (UNK1) | 118 | 2021 | 539 | 800 | 582 | 4060 |
| Unknown 2 (UNK2) | 239 | 804 | 38 | 42 | 24 | 1147 |
| Fibroblast (FB) | 116 | 589 | 92 | 194 | 127 | 1118 |
| Mesangial (MES) | 132 | 244 | 47 | 189 | 40 | 652 |
| Myocytes | 0 | 21 | 13 | 41 | 11 | 86 |
| **Total per condition** | **12961** | **27814** | **5536** | **12338** | **5829** | **64478** |

nNK: native normal kidney, NKTx: normal allograft, nFSGS: native kidney with FSGS, KTxFSGS-T1: kidney allograft with first recurrence of FSGS post-kidney transplantation, KTxFSGS-T2: kidney allograft with second recurrence of FSGS post-kidney transplantation.

**Table S4: Top markers for cell clusters**

| **Cell Cluster** | **Gene markers** |
|---|---|
| Proximal tubule 1 (PT-1) | *CUBN, AFM, SLC36A2, PLG, PAH, SLC13A1, AGXT2, SLC34A1, SMIM2-AS1, SLC13A3, ACSM2B, ACSM2A, CLDN10, AGXT2* |
| Distal tubule (DT-1) | *SLC12A1, CLCNKB, MSFD4A, COL28A1* |
| Endothelial (EC-1, EC-2, EC-3) | *IGFBP5, HEG1, PECAM1, EGFL7, PTPRB, NOTCH4* |
| Distal tubule 2 (DT-2) | *THRB, CLCNKB, WNK4, TMPRSS2, DEFB1, CACNA1D, PIK3C2G, TFCP2L1, SGMS2, SLC8A1* |
| Collecting duct principal 1 (CDP-1) | *CADPS2, ATP1B1, NRDG1, SNTG1, CALB1, SCNN1G, SCN2A* |
| Proximal tubule 2 (PT-2) | *HAVCR1, DCC, DLGAP1, MYO3A, LINC02471, CLSTN2, VCAM1, ACSM2A, SLC47A2, SLC17A1, GLYAT, MYO7B* |
| Collecting duct principal 2 (CDP-2) | *AQP2, AQP3, COBLL1, MECOM3, FXYD4* |
| Collecting duct intercalated 1 (CDI-1) | *ADGRF5, ADGRF1, AQP6, DMRT2, ATP6VOD2* |
| Podocyte (POD) | *WT1, NPHS2, NHPS1, ADAMTS19, FMN2, NTNG1, PTPRQ, ST6GALNAC3* |
| Parietal Epithelial (PEC) | *KIRREL3, SLC4A11, LINC01435, CFH, KLRG2, ALDH1A2, KCNT2, CLDN1, RBFOX1*, *PAX8* |
| Immune (IMM) | *IKZF1, ITGAL, PIK3R5, ITGA4, CD96, CD44* |
| Collecting duct intercalated 2 (CDI-2) | *INSRR, RHBG, SLC26A4, PD2D2, TMEM213, SCIN, FOXI1, WFDC2* |
| Fibroblast (FB) | *FBLN1, ADH1B, DCN, C7, PDGFRA, SLIT2* |
| Mesangial (MES) | *AGTR1, PIEZO2, DAAM2, CARMN, PRR16, ITGA8, COL25A1, ROBO1* |
| Myocytes | *PRKG1, SYNPO2, NEXN, CAP2, TTN, DMD* |

ADDITIONAL SUPPLEMENTARY TABLES ARE PRESENT IN THE ‘Supplementary Tables.xlsx’ FILE

**Fig. S1: Quality control parameters of the study.** These included (A) percent mitochondrial gene expression per cell (percent.mito) versus the total number of RNA detected per cell (nCount_RNA), (B) the number of genes detected in cell (nFeature_RNA) versus nCount_RNA, also depicted are (C) nFeature_RNA, nCount_RNA, and percent.mito per sample. The cut off used in the study is present in the main text.

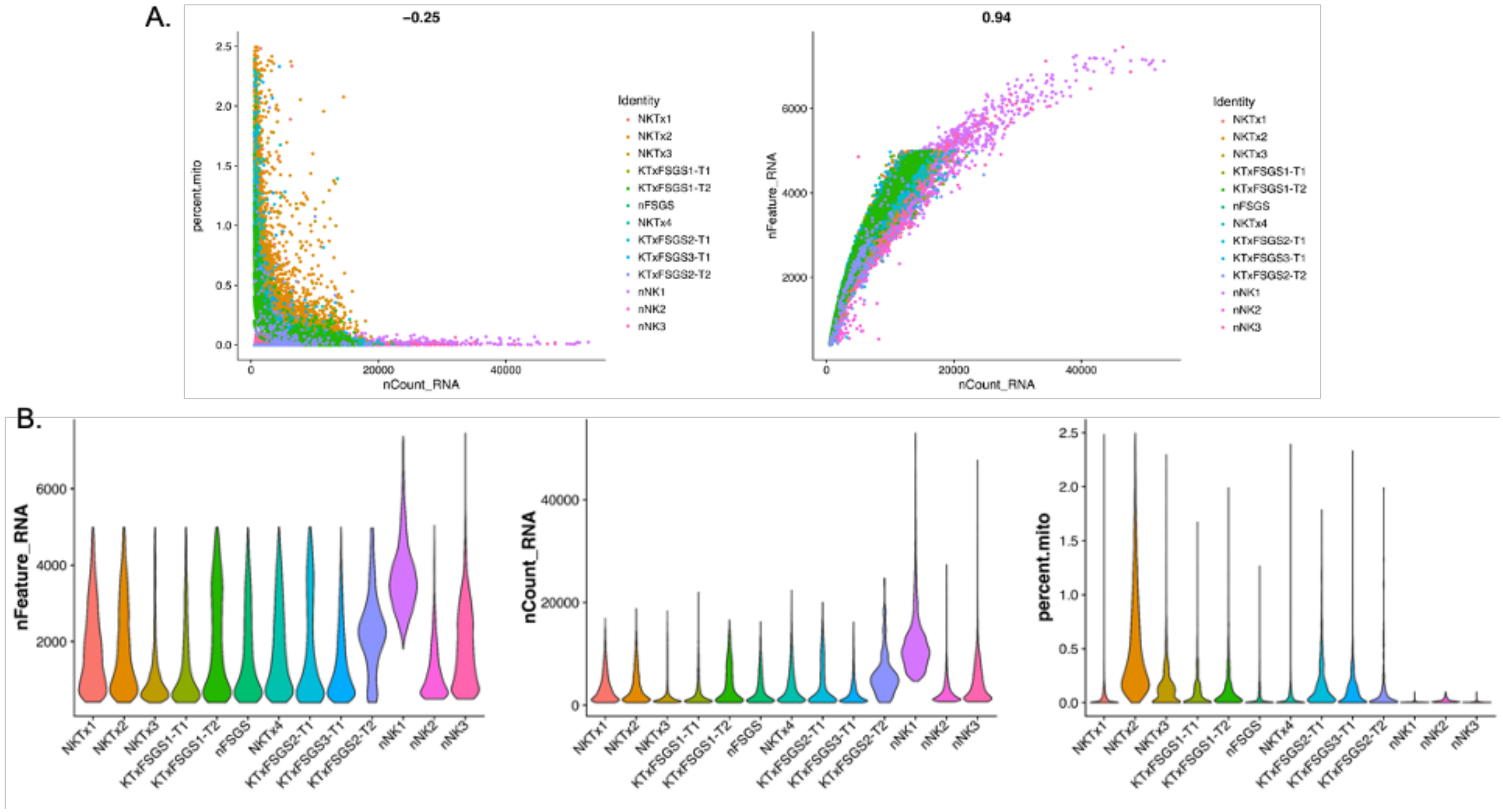

**Fig. S2: Cell cluster identification.** (A) Total cell clusters identified in our single nuclei (sn) RNA sequencing presented as an integrated uniform manifold approximation and projection (UMAP). (B) The integrated UMAP was split based on patient condition consisting of normal native kidney - nNK, kidney allograft with normal function - NKTx, native kidney with focal segmental glomerulosclerosis (FSGS) - nFSGS, transplanted kidney with recurrent FSGS at first collection – KTxFSGS-T1, and second collection during the next recurrence - KTxFSGS-T2. Nineteen clusters of cells including the two clusters that had markers of multiple cell types (UNK-1 and UNK-2) which were excluded from the study in our current downstream analysis.

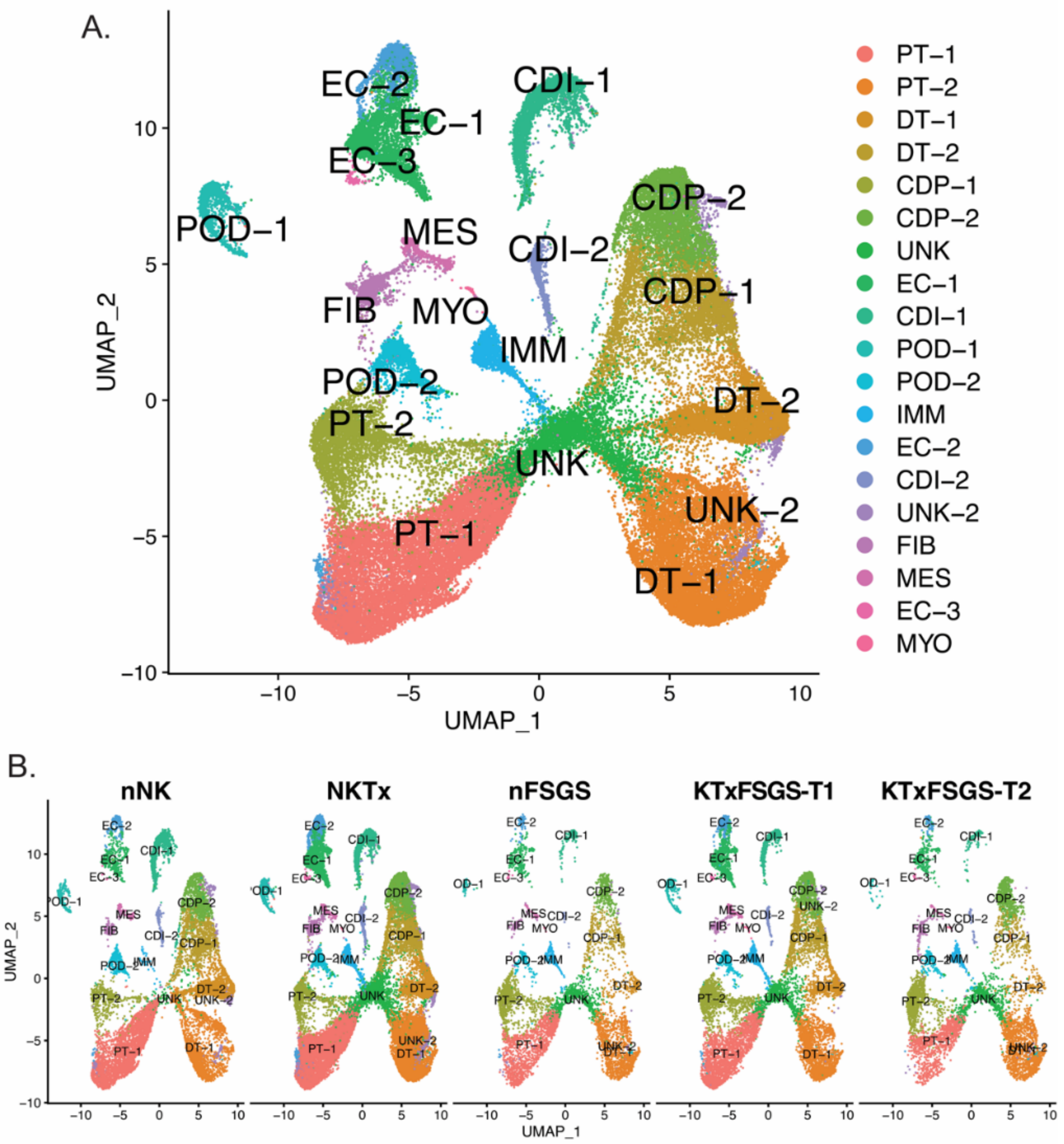

**Fig. S3: Treatment of conditionally immortalized podocytes with serum of patients with recurrent FSGS.** Podocytes were grown under immortalization conditions as per the vendor's instructions after which they were allowed to differentiate for 5 days in the differentiation media. On the 5th day, the cells were cultured with media supplemented with 4% of patient serum (N = 2) and cultured for additional 48 hours after which the RNA was collected and Taqman based qPCR (n=2) was used to determine expression of genes mentioned in the figure, p-value * < 0.05 (Student's t-test).

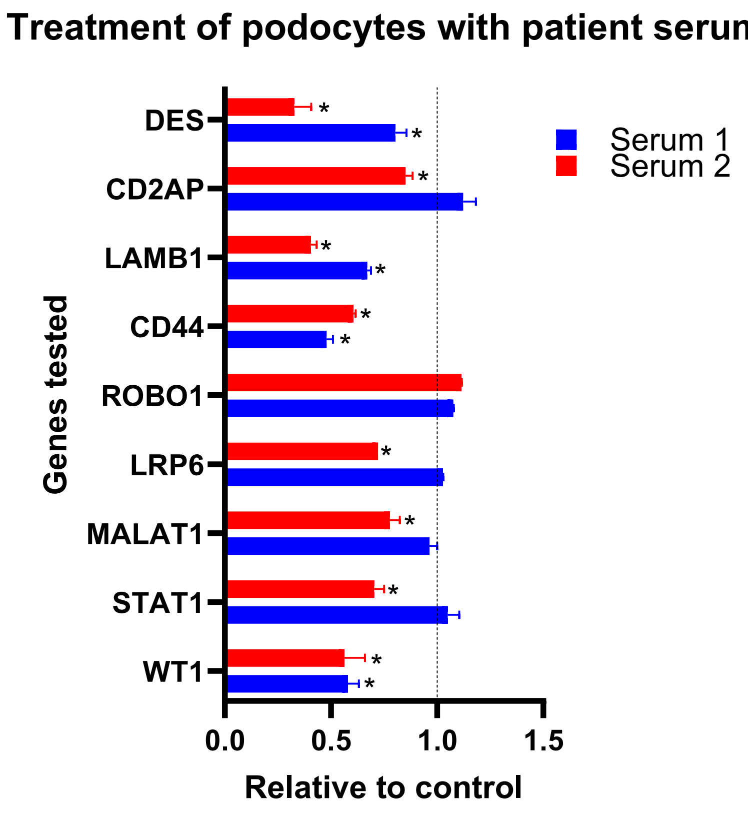

**Fig. S4: Biological properties associated with differentially expressed genes between NKTx and KTxFSGS.** (A) EnrichR based identification of pathways associated with significantly downregulated genes in podocytes of KTxFSGS-T1 compared to KTx. (B)DotPlot of cell death related genes in NKTx (Normal) and post-transplant recurrent FSGS (PostTXwFSGS) does not demonstrate an enrichment of apoptosis-associated genes in either condition.

(A)

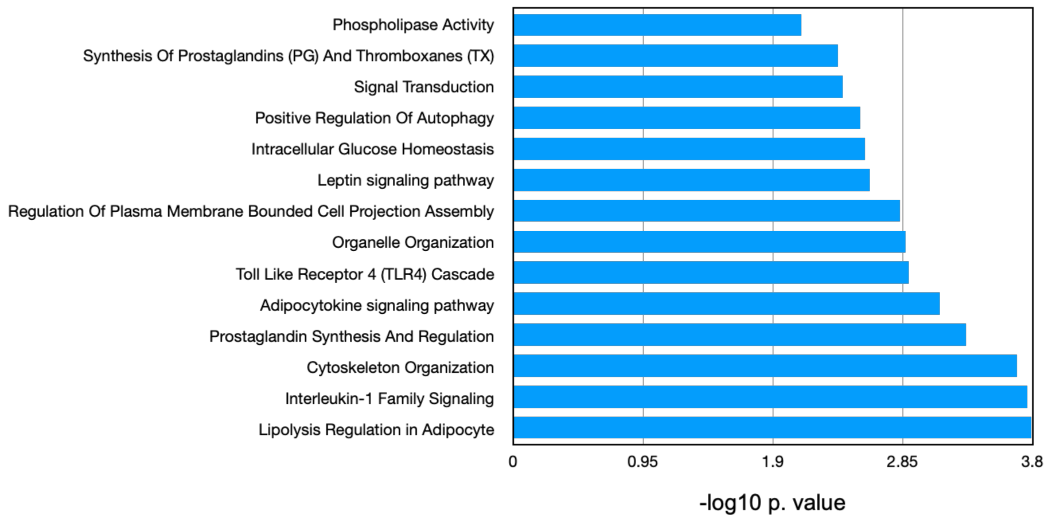


(B)

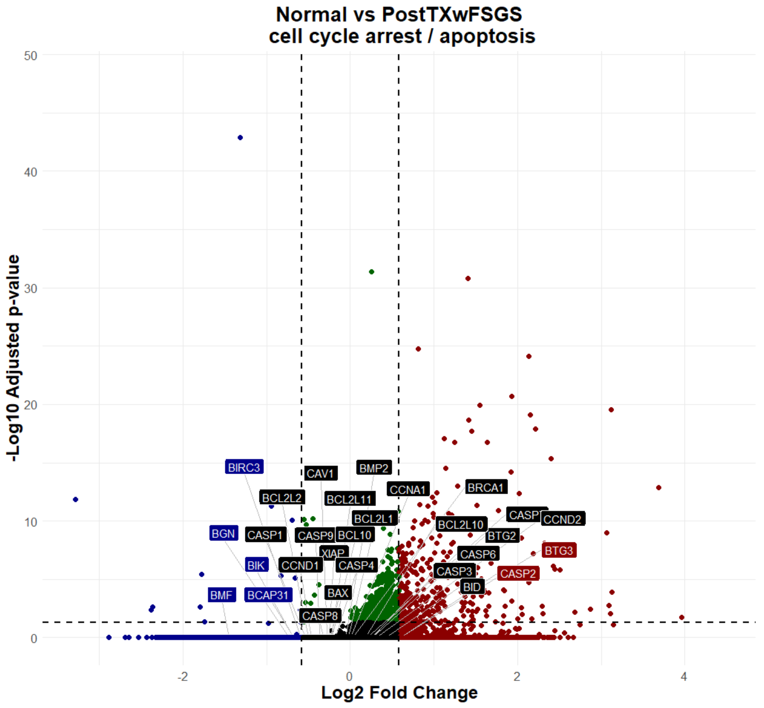

**Fig. S5:** Metascape-based pathway analysis of differentially expressed genes in podocytes of individual recurrent T1-FSGS patient samples (FSGS1-3) compared to normal allograft (NKTx)

(A) Downregulated genes in FSGS1 (v NKTx)

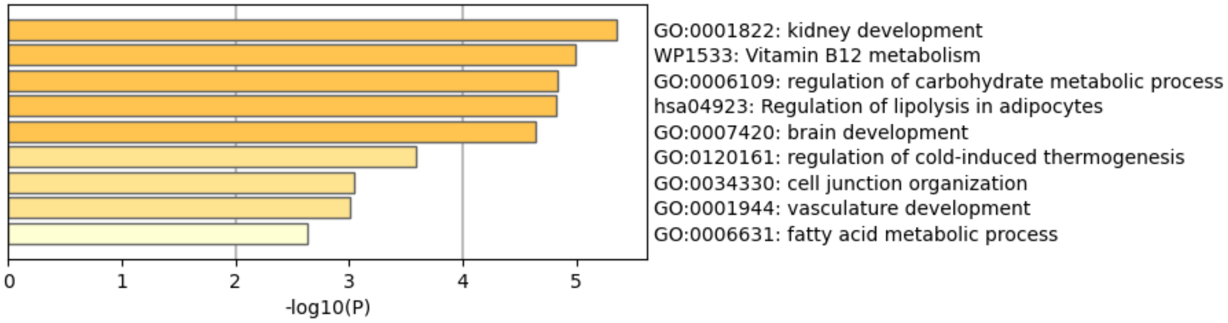


Upregulated genes in FSGS1 (v NKTx)

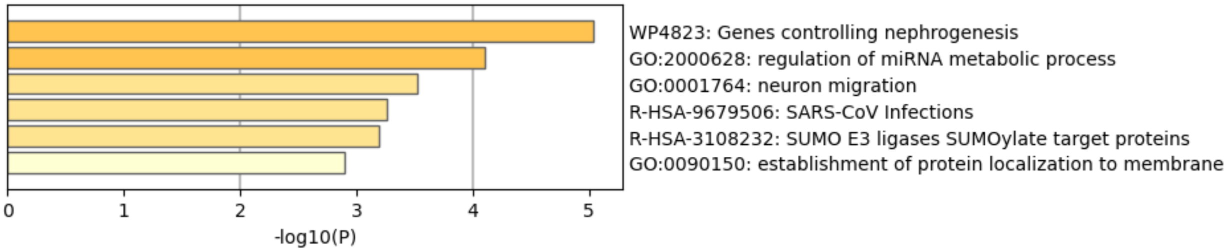


(B) Downregulated genes in FSGS3 (v NKTx)

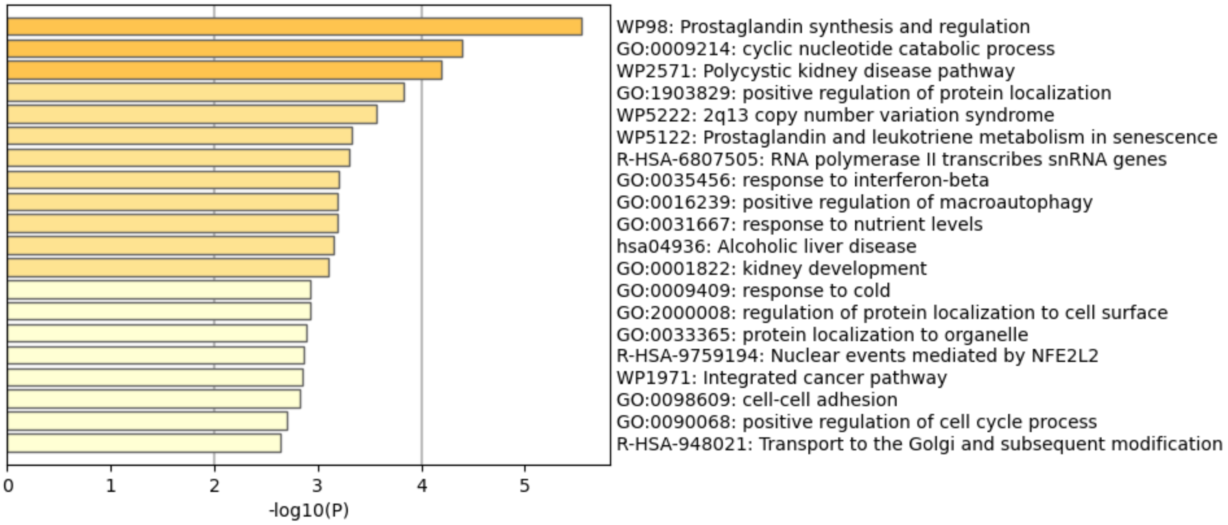

**Fig S6:** DotPlot demonstrating the presence of CD44 in podocytes cluster (POD) of FSGS2 (POD_GEX-NW-5). Classical podocyte associated genes have also been plotted for reference.

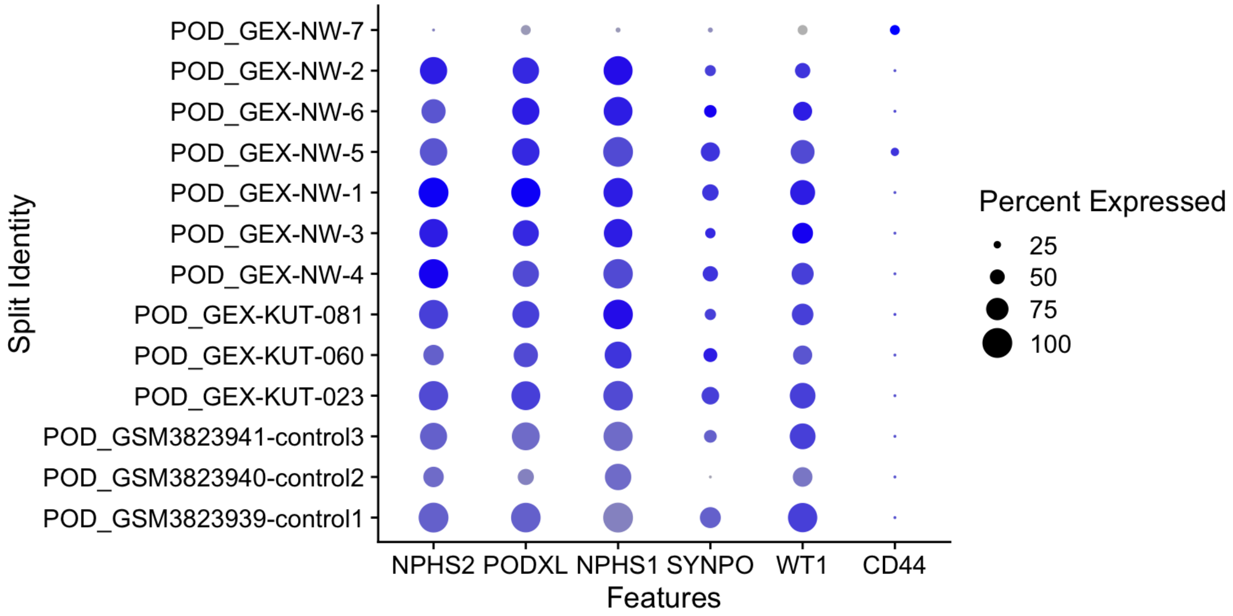


(<u>CD44 alone</u>)

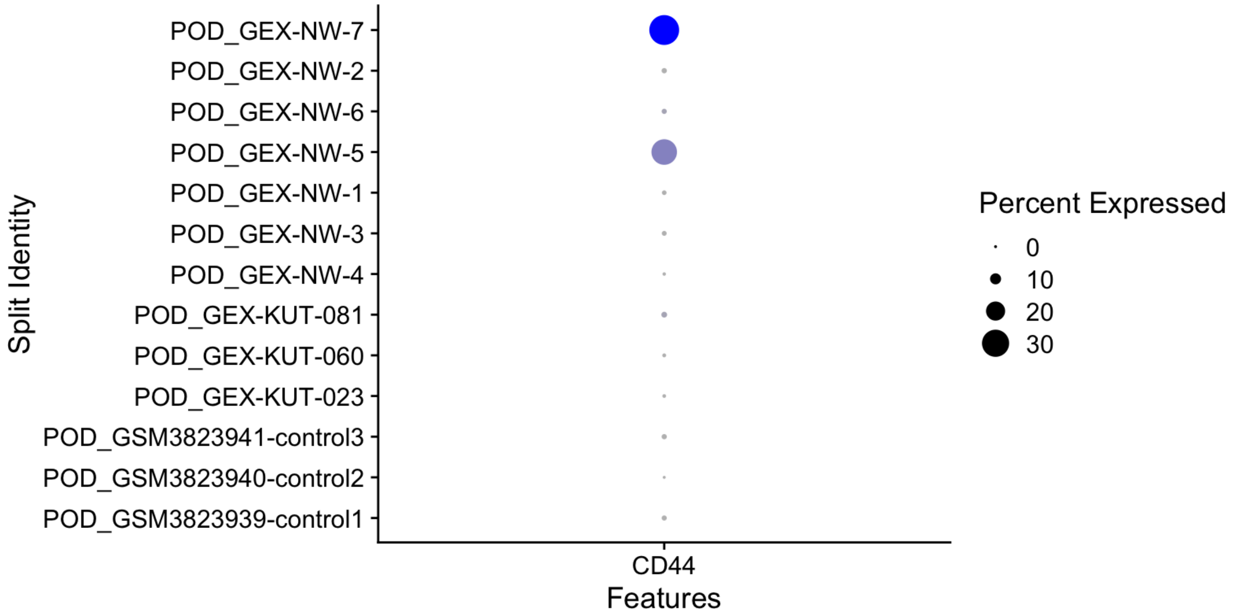

**Fig. S7:** DotPlot demonstrating the presence of CD44, HB-EGF, and EGF in PEC cluster, represented in (A) all conditions, and (B) all normal allograft samples.

**(A)**

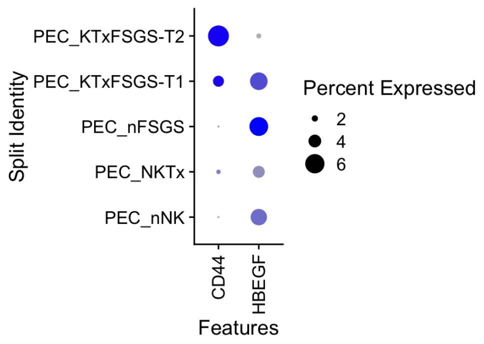


**(B)**

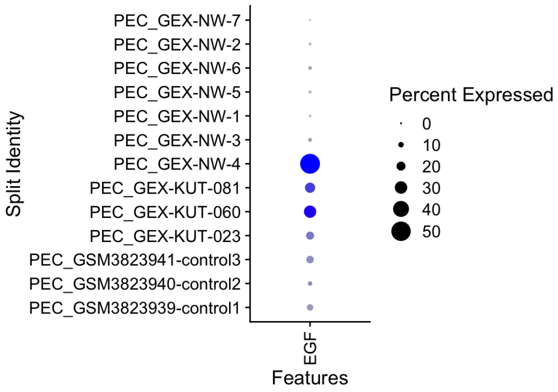

**Fig. S8:** Metascape-based pathway analysis of differentially expressed genes glomerular endothelial cells.

A. Upregulated genes in nFSGS (v nNK)

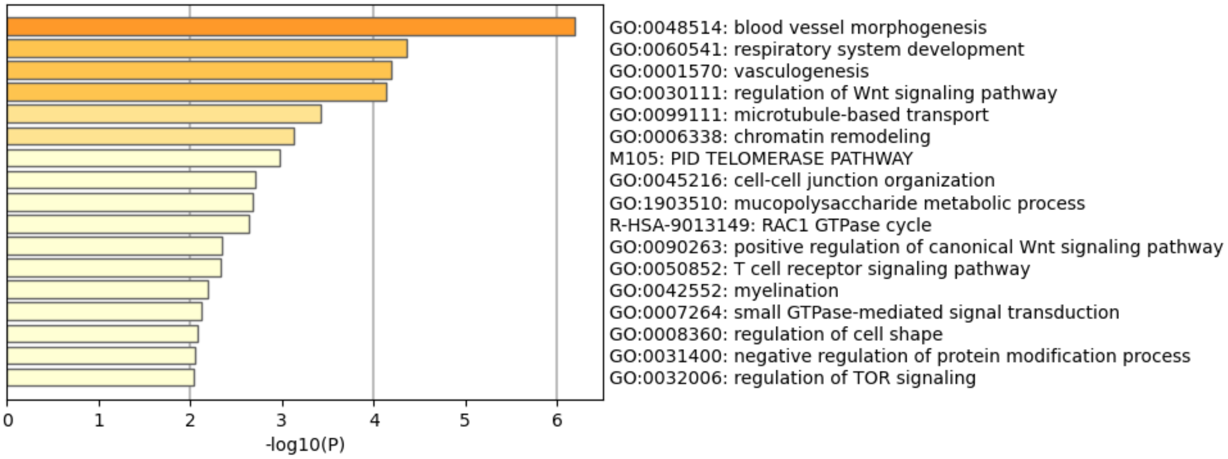


B. Downregulated genes in nFSGS (v nNK)

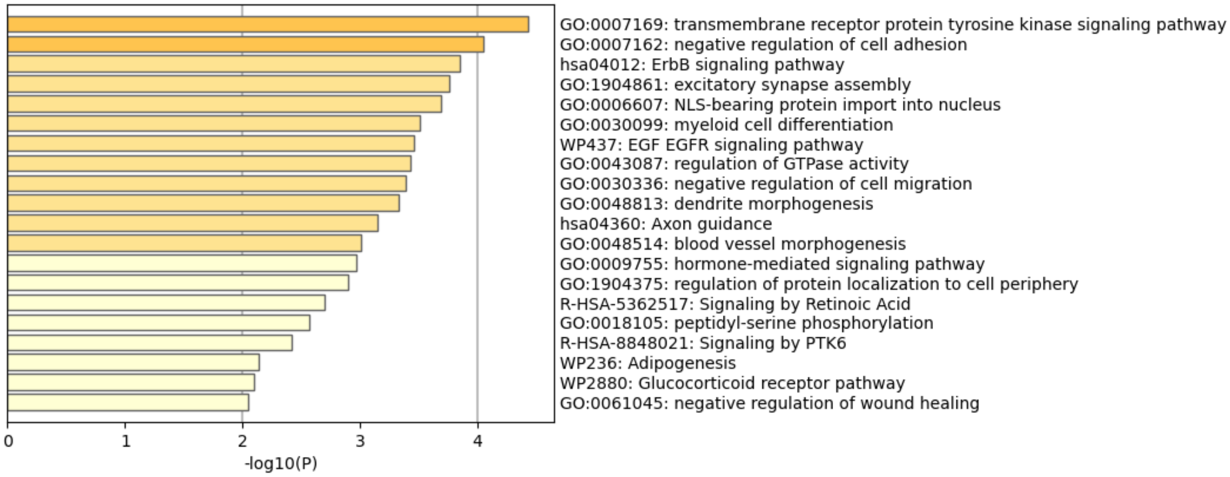

**Fig. S9:** Metascape-based pathway analysis of differentially expressed genes of proximal tubular cells of nFSGS (v nNK).

A. Dot plot demonstrating expression of injury markers and progenitor markers in FSGS samples

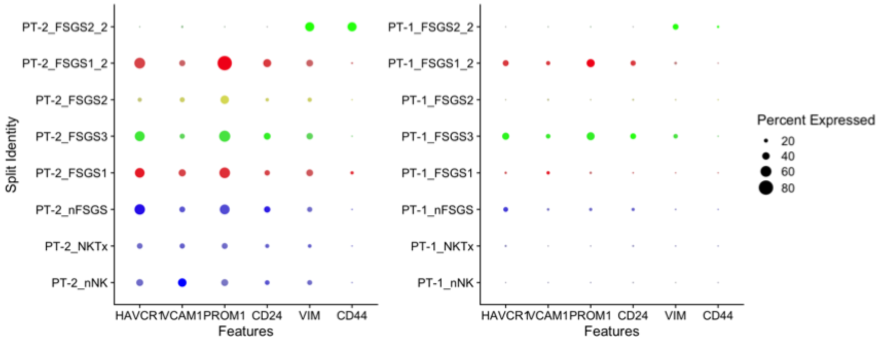


B. Pathway analysis of cluster PT-1 associated differentially expressed genes

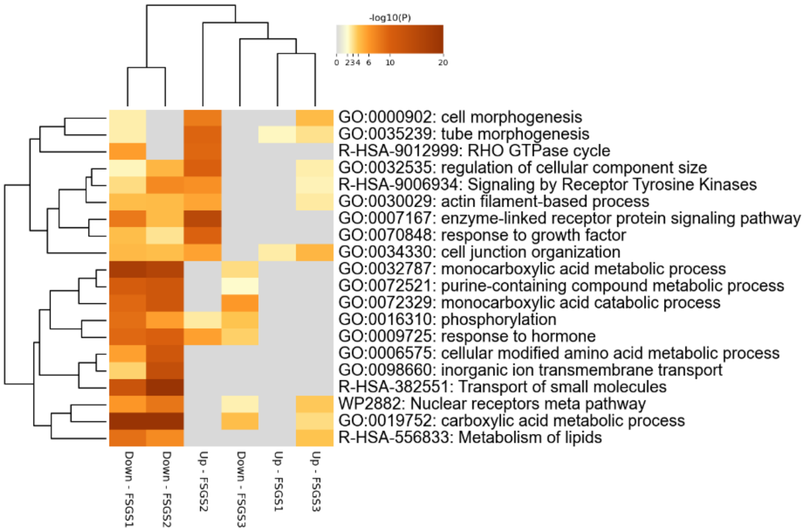


C. Pathway analysis of cluster PT-2 associated differentially expressed genes

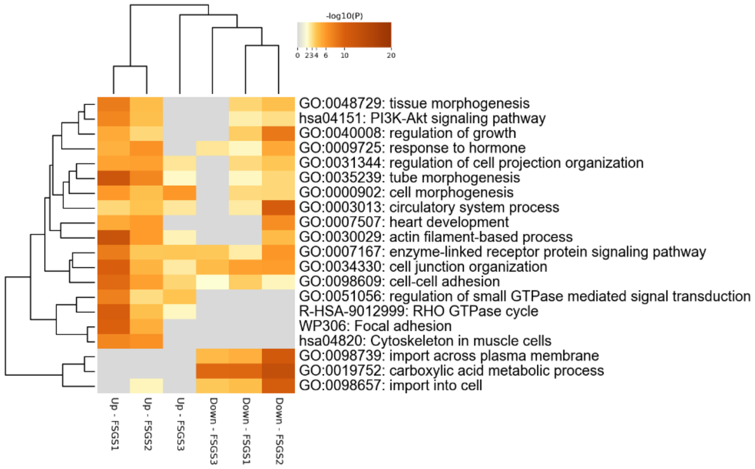
